# Multi-physics simulations reveal hemodynamic impacts of patient-derived fibrosis-related changes in left atrial tissue mechanics

**DOI:** 10.1101/2024.05.29.596526

**Authors:** Alejandro Gonzalo, Christoph M. Augustin, Savannah F. Bifulco, Åshild Telle, Yaacoub Chahine, Ahmad Kassar, Manuel Guerrero-Hurtado, Eduardo Durán, Pablo Martínez-Legazpi, Oscar Flores, Javier Bermejo, Gernot Plank, Nazem Akoum, Patrick M. Boyle, Juan C. del Alamo

## Abstract

Stroke is a leading cause of death and disability worldwide. Atrial myopathy, including fibrosis, is associated with an increased risk of ischemic stroke, but the mechanisms underlying this association are poorly understood. Fibrosis modifies myocardial structure, impairing electrical propagation and tissue biomechanics, and creating stagnant flow regions where clots could form. Fibrosis can be mapped non-invasively using late gadolinium enhancement magnetic resonance imaging (LGE-MRI). However, fibrosis maps are not currently incorporated into stroke risk calculations or computational electro-mechano-fluidic models. We present multi-physics simulations of left atrial (LA) myocardial motion and hemodynamics using patient-specific anatomies and fibrotic maps from LGE-MRI. We modify tissue stiffness and active tension generation in fibrotic regions and investigate how these changes affect LA flow for different fibrotic burdens. We find that fibrotic regions and, to a lesser extent, non-fibrotic regions experience reduced myocardial strain, resulting in decreased LA emptying fraction consistent with clinical observations. Both fibrotic tissue stiffening and hypocontractility independently reduce LA function, but together, these two alterations cause more pronounced effects than either one alone. Fibrosis significantly alters flow patterns throughout the atrial chamber, and particularly, the filling and emptying jets of the left atrial appendage (LAA). The effects of fibrosis in LA flow are largely captured by the concomitant changes in LA emptying fraction except inside the LAA, where a multi-factorial behavior is observed. This work illustrates how high-fidelity, multi-physics models can be used to study thrombogenesis mechanisms in patient-specific anatomies, shedding light onto the links between atrial fibrosis and ischemic stroke.

**Key points:**

- Left atrial (LA) fibrosis is associated with arrhythmogenesis and increased risk of ischemic stroke; its extent and pattern can be quantified on a patient-specific basis using late gadolinium enhancement magnetic resonance imaging.
- Current stroke risk prediction tools have limited personalization, and their accuracy could be improved by incorporating patient-specific information like fibrotic maps and hemodynamic patterns.
- We present the first electro-mechano-fluidic multi-physics computational simulations of LA flow, including fibrosis and anatomies from medical imaging.
- Mechanical changes in fibrotic tissue <u>impair</u> global LA motion, decreasing LA and left atrial appendage (LAA) emptying fractions, especially in subjects with higher fibrosis burdens.
- Fibrotic-mediated LA motion impairment alters LA and LAA flow near the endocardium and the whole cavity, ultimately leading to more stagnant blood regions in the LAA.

## 1. Introduction

Stroke is a leading cause of mortality and disability worldwide (1). About 75% of all strokes are ischemic strokes (1), in which blood flow to the brain is reduced or blocked. Ischemic stroke risk increases 5-fold in patients with atrial fibrillation (AFib) (2), a common arrhythmia affecting about 38 million worldwide (3). During AFib, the unsynchronized motion of the atrial myocardium causes blood stagnation, promoting thrombogenesis. Atrial thrombi generally form in the left atrial appendage (LAA) – a small, narrow enclosure extending off the left atrium (LA). Ischemic strokes produced by embolization of these clots are associated with more disability and higher mortality than ischemic strokes not due to this mechanism, such as those due to atherosclerotic vascular disease (4, 5).

AFib patients are treated with anticoagulation medication to reduce their thrombotic risk, but the treatment increases their bleeding risk (6). The current tools to assess whether anticoagulation therapy is warranted for a particular patient (e.g., the CHA_2_DS_2_VASc score) are based on comorbid factors, which lack any mechanistic associations with thrombus formation, predictive accuracy, and personalization (7). Notably, the same metrics used to calculate CHA_2_DS_2_VASc are associated with fibrotic myocardial remodeling in response to tissue injury (8). The “C” in CHA_2_DS_2_VASc indicates congestive heart failure, which can be attributed to the development of fibrous connective tissue (9). Fibrosis is associated with AFib and is higher in patients with a history of stroke (“S_2_”) than in those without reported strokes (10). Fibrosis is also known to increase with age (“A_2_” and “A”) (11). Finally, women with AFib and stroke have a higher fibrosis burden than men (“Sc”) or other women with AFib but without stroke history (12).

Despite the broad medical evidence associating fibrosis with AFib and stroke (10, 13, 14), fibrosis is not included in risk stratification tools. While this exclusion is partly due to fibrotic mapping requiring non-routine imaging testing, it can also be attributed to our currently limited understanding of the mechanisms by which fibrosis may promote atrial thrombosis. In fibrotic regions, replacement of dead cardiomyocytes or overproduction of extracellular matrix alters the electrical conduction velocity and the biomechanical properties of the myocardium (15). In particular, the tissue becomes stiffer and hypocontractile (15, 16). These changes are recognized to reduce myocardial kinetics, causing areas of slow flow that can increase the risk of thrombosis (14, 17). However, each factor’s contribution, both in isolation and interacting with other factors, is poorly understood. We also lack knowledge about how fibrotic derangement of myocardial kinetics disturbs atrial hemodynamics.

There is a great clinical need for a personalized approach in the management of AFib, especially in stroke prediction and prevention, routed in pathophysiological mechanisms that explain associations demonstrated in epidemiological studies (18). Advanced imaging modalities that allow for non-invasive, detailed quantification of fibrotic remodeling motivate the conduction of mechanistic studies of the nexus between AFib, atrial fibrosis, and the hemodynamic thrombotic substrate. Multi-physics computational models are ideally suited for these investigations since they permit manipulations that are not feasible in animal or clinical experiments. These models also produce high-resolution spatiotemporal data that facilitate in-depth quantitative analyses (17). Numerous research groups have developed 3D-coupled electromechanical and hemodynamics computational frameworks. Different models are used to solve the systems of mathematical equations that represent electric potential propagation, active/passive biomechanics, and fluid dynamics; the schemes used to couple these different systems (i.e., loose/weak or full/strong) are also variable. Likewise, idealized, realistic, and imaged-based anatomy models of the chambers and valves have been used in different studies. For detailed reviews on this topic, the reader is referred to (19, 20).

Watanabe et al. (21) pioneered 3D multi-scale, multi-physics models in left ventricular (LV) geometries, with simulations producing physiological pressure values and aortic flow rates. The most recent version of this solver, the monolithic UT-Heart (22), has been used to simulate surgical planning for cardiac resynchronization therapy. Vigmond et al. (23) performed the first whole heart simulations in an idealized geometry to investigate the impact of bundle branch block on transvalvular flow, septal movement, and stretch-induced effects for myocytes in the left and right ventricular walls. Quarteroni et al. (24) introduced multi-physics computational models of idealized LVs, which were subsequently improved to consider realistic left heart (25) and whole-heart anatomies (26). Feng et al. (27) analyzed the effects of mitral valve regurgitation and AFib on LA and LAA flow in an LA-mitral valve system using their coupled fluid interaction model. Viola et al. (28) introduced a fluid-structure-electrophysiology interaction solver in an anatomy built from a medical atlas, and Bucelli et al. (25) performed simulations in a realistic average left heart model. Zingaro et al. (26) built on that framework to include flow through the coronary arteries and a myocardial perfusion model. Choi et al. (29) introduced their methodology combining MRI-based electromechanics and a hemodynamics solver to investigate flow differences in normal and failing canine hearts. Most recently, Viola et al. (30, 31) presented a digital twin model of the human heart to simulate left bundle branch disorders.

Despite these significant advances in multi-physics cardiac computational models, only one simulation study has incorporated fibrotic effects on intracardiac flow: Paliwal et al. (32) used computational fluid dynamics (CFD) to investigate aberrant hemodynamic patterns in subjects with a higher fibrosis burden and higher values of pro-thrombotic hemodynamic metrics near fibrotic regions. These simulations used an idealized LA wall motion based on the change of LA volume over time instead of resolving the intricate nature of fibrosis and its effects on myocardial biomechanics.

This study presents a multi-physics, multi-scale computational framework with coupled models of EP, biomechanics, and CFD. We used this framework to study the effects of fibrosis on LA hemodynamics. We simulated LA contraction in four subjects with different burdens of fibrotic remodeling, focusing on the resulting biomechanical derangement. In our simulations, higher fibrosis burdens cause more severe impairments of atrial motion, leading to decreased LA and LAA emptying fractions (EF). Decreased EF is linked to smaller kinetic energies in the LAA that could predispose patients to thrombosis. We present a methodology to better understand the mechanisms underlying these disturbances by simulating the isolated and combined impacts of fibrotic tissue stiffening and loss of contractility, as well as by digitally restricting fibrotic patches to specific atrial regions. The tools and approaches presented here constitute a significant step towards building left atrial models with patient-specific anatomy and distribution of fibrosis. Multi-physics simulations of these models enable unprecedented physiological thought experiments and hold potential to support clinical decisions and investigate therapeutic targets.

## 2. Methods

This section describes the patients analyzed, the image acquisition protocol, the meshing approach, the multi-physics, the multi-scale computational framework (including our approach to representing passive and active biomechanical consequences of fibrosis), and the analysis methods to quantify atrial wall motion and blood flow.

### 2.1. Patients studied, image acquisition, and mesh generation

As the starting point for our analysis, we retrospectively used LGE-MRI scans of the LA from four patients, all of which were collected for clinical use (33-35). Approval from the institutional review board (IRB #8763) within the University of Washington Cardiac Arrhythmia Data Repository (CADRE database) was obtained. Prior to enrollment, study participants provided informed consent, and the investigation adhered to the principles outlined in the Declaration of Helsinki. None of the patients had prior catheter ablation, and therefore, the fibrosis quantified was not due to ablation-induced scarring. Demographics and comorbidity information obtained at the time of the MRI scans for all four patients are shown in Table 1. The rationale for choosing these four particular cases was to obtain two representative cases each of low (7% and 12%) and high fibrosis (41% and 47%) burdens, as assessed by total volume of fibrotic myocardium in our volumetric 3D reconstructions (see detailed methodology below). The naming scheme for models uses the format “FibNN”, where NN is the percentage of fibrotic tissue volume. The method used for LGE-MRI scan acquisition has been described extensively in our prior work (36-38). Briefly, studies were performed on a Philips Ingenia 1.5T scanner; images were obtained 15-25 minutes after delivery of gadolinium contrast; a 3D inversion recovery, respiration navigated, ECG-gated, gradient echo pulse sequence was used to acquire a transverse imaging volume with voxel size = 1.25×1.25×2.5 mm_3_ (reconstructed to 0.625×0.625×1.25 mm_3_). LGE-MRI analysis of fibrosis was performed by the Merisight service (Marrek Inc., Salt Lake City, UT); the result of this process was a 3D surface geometry of the LA annotated with values indicating the presence or absence of fibrosis.

**Table 1.** Subjects’ demographics, comorbidities, and fibrotic information. For each subject, we provide the CHA2DS2VASc score and the contributing factors: age, sex, history of congestive heart failure, stroke, coronary artery disease, hypertension, and diabetes. LA fibrosis was obtained as the total volume of fibrotic myocardium tissue detected by LGE-MRI acquisitions.

|  | **Fib07** | **Fib12** | **Fib41** | **Fib47** |
| --- | --- | --- | --- | --- |
| **Age (years)** | 51 | 60 | 67 | 55 |
| **Sex** | Female | Female | Male | Female |
| **Congestive heart failure** | No | Yes | No | Yes |
| **Hypertension** | Yes | No | Yes | No |
| **Diabetes mellitus** | No | No | No | No |
| **Stroke** | Yes | No | No | Yes |
| **Coronary artery disease** | No | No | No | No |
| **CHA<sub>2</sub>DS<sub>2</sub>VASc score** | 3 | 2 | 2 | 3 |
| **LA fibrosis (%)** | 7.0 | 12.0 | 41.1 | 47.1 |

#### 2.1.1. Volumetric mesh generation

From the surfaces generated by Merisight, we constructed 3D volumetric meshes using *meshtool (39)*, an open-source finite element meshing and data manipulation software (available: https://bitbucket.org/aneic/meshtool). First, we resampled the surface meshes to an average edge length of approximately 0.8 mm, sufficient to resolve the anatomical details of left atrial geometries under consideration. Next, we extruded the resampled surface meshes – corresponding to the interface between the blood pool and the tissue – with four layers, each having a thickness of 0.5 mm. This process resulted in an overall uniform LA wall thickness of 2 mm. The deliberate choice of a smaller edge length in the transmural direction aimed to mitigate numerical issues, such as locking effects commonly encountered in tissue mechanics simulations involving thin-walled geometries. The extrusion produced four layers of prisms, which were subsequently converted into tetrahedral elements for the finite element computations.

We used the method described by Roney et al. (40) to assign endocardial and epicardial fibers. Subsequently, we interpolated element tags from the original surface meshes – labeling healthy and fibrotic tissue – onto the resampled surface meshes and propagated them through the tissue in a transmural direction.

### 2.2. Multi-scale, multi-physics computational framework

#### 2.2.1. Multi-scale electromechanical modeling

Myocardial electromechanics (EM) was modeled on patient-specific anatomies and fibrotic maps using the in-house code Carpentry (41); a version of this software capable of carrying out electrophysiological (EP) simulations (openCARP) is publicly available for non-commercial use (42). To represent human atrial action potentials, we used the cellular EP model by Courtemanche et al. (43) with modifications to represent changes in patients with AF (44). This is recognized as a standard approach for representing cell-scale electrophysiology in the diseased atria (45). The calculation of intercellular current flow, responsible for propagating electrical activation in the atrial myocardium, was performed using the monodomain equation. The decision to use this formalism instead of the more mathematically burdensome bidomain equations is justified, since the models in this study do not necessitate accurate representation of current injection into the extracellular space (46). For healthy tissue, conductivities in the longitudinal, transverse, and sheet normal directions were chosen as *g*_*l*_ = 10.551 S/m and *g*_*t*_ = *g*_*n*_ = 0.086 S/m, respectively. To account for a slower conduction velocity in fibrotic tissue, these values were changed to 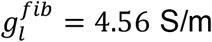 and 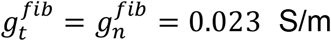 in fibrotic regions. These values were chosen to comport with the longitudinal:transverse ratios used in our previous studies (2.3:1 non-fibrotic; 3.7:1 fibrotic) (33-35). Isotropic extracellular conductivity was uniformly set to *g*_e_ = 0.8 S/m in all regions, and the harmonic mean of intracellular and extracellular conductivities produced effective monodomain conductivity values corresponding to negligible conduction velocity (CV) slowing in the longitudinal direction (∼5%) and modest CV slowing in the transverse direction (∼46.3%). During normal sinus rhythm, observations from electroanatomic mapping (35) suggest that LA excitation originates from a small number of points where bundles originating in the right atrium (including Bachmann’s bundle) become contiguous with the LA. Accordingly, excitation in our models was initiated via transmembrane current stimulation in a prescribed region corresponding to the typical penetration site of Bachmann’s bundle on the anterior wall. Wavefronts then propagated in an anisotropic, heterogeneous manner governed by the underlying conductivity tensors used to represent the fibrous structure of the LA. **Figure 1A** shows examples in a simple tissue geometry for these nominal CV values under non-fibrotic and fibrotic conditions. In the current modeling framework, we excluded cell-scale EP effects of fibrosis since they affect the shape of the intracellular calcium transient and impair contraction, hindering the assessment of biomechanical derangements in fibrotic regions.

**Figure 1.**
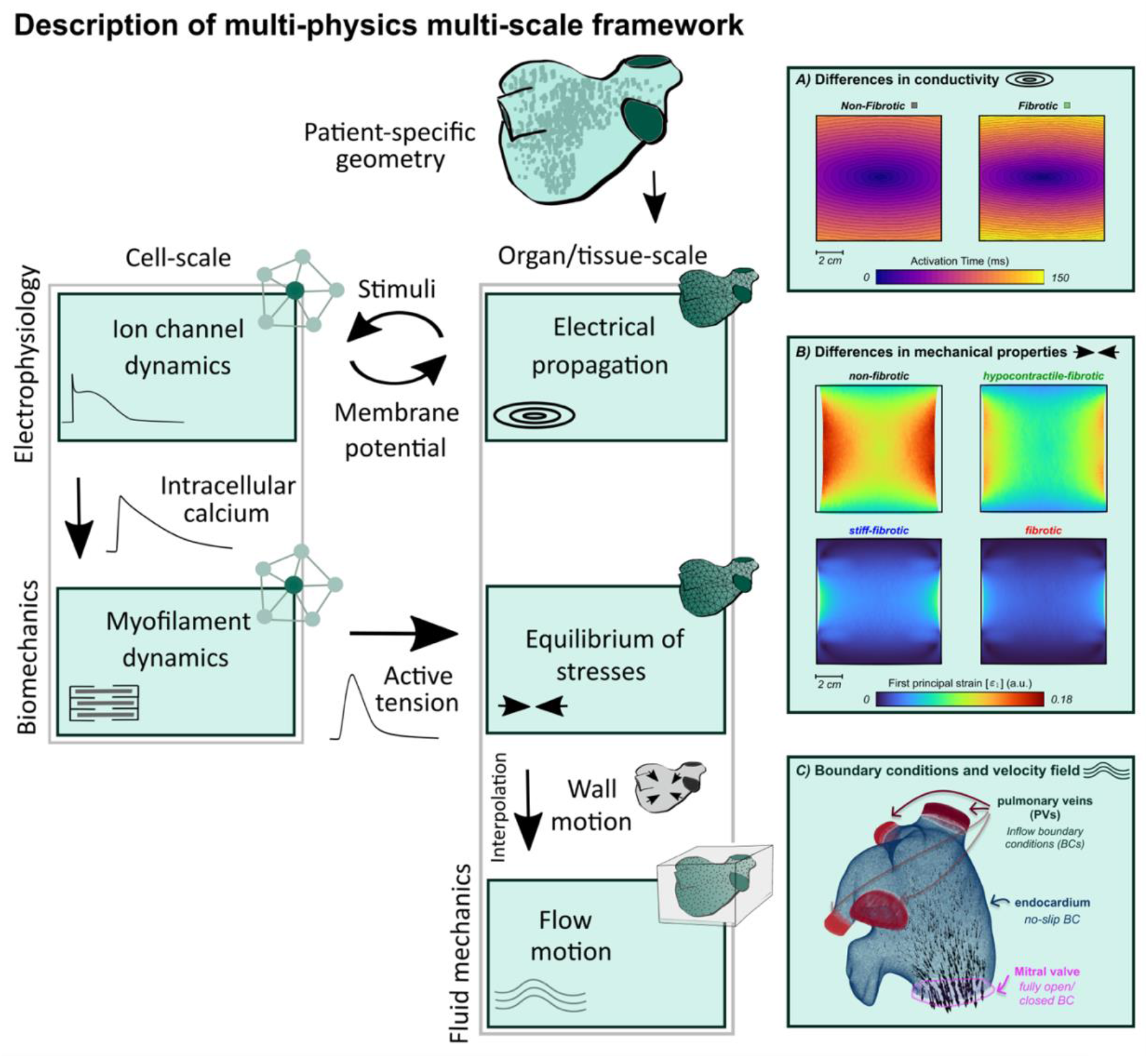
Description of multi-physics multi-scale framework. Block diagram of electrophysiology and biomechanics multi-scale models, and fluid mechanics model including coupling between them. **A)** Differences in fibrotic and non-fibrotic tissue conductivity depicted with activation time maps in an anisotropic 10 cm2 tissue patch highlighting impaired transverse conduction. **B)** Differences between non-fibrotic, hypercontractile-fibrotic, stiff-fibrotic, and fibrotic mechanical states shown with *ε*_1_ maps in the same tissue patch. **C)** Sketch of computational fluid dynamics model approach to impose boundary conditions including velocity field vectors obtained during LA systole in black.

The passive tissue of the myocardium of the LA was modeled as a transversely isotropic material (see Augustin et al. 2020 (47)) with the strain energy function

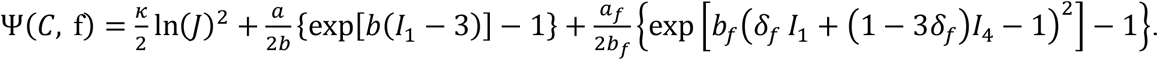

Here, the first term of the equation is the volumetric contribution to ensure nearly incompressible material behavior, *κ* = 650 kPa is the bulk modulus, and *J* is the determinant of the deformation gradient F. The second term models isotropic tissue components using constants *a* = 2.92 kPa and *b* = 5.6 and, and the first invariant 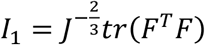. The third term is the anisotropic contribution in myocyte fiber direction *f* modeled using constants *a*_*f*_ = 11.84 kPa, *b*_*f*_ = 17.95, and *δ*_*f*_ = 0.09, and the invariant *I*_4_ = *f* · *F*^*T*^*Ff*. For fibrotic tissue, we multiplied the stress-dependent parameters *a* and *a*_*f*_ by a fixed factor of five, based on limited experimental data (48, 49).

For each of the four geometries we performed a backward displacement algorithm (50) to find an unloaded reference configuration for the healthy and fibrotic case and then in a second step reloaded the reference geometries using a given prescribed pressure. This unloading-reloading procedure ensures alignment between the pressurized healthy and fibrotic models and the configuration obtained from image data. As the exact pressure at the time of imaging in the LA was unknown, we adopted a prescribed pressure of 10 mmHg as in (47).

We subjected these pressurized geometries to an active stress experiment where the pressure remained constant throughout the simulation (see section 2.3 for more details). The active stress tensor resulting cardiac myocytes contraction was assumed to exert force in the myofiber direction f, where we also consider a certain amount of fiber dispersion following the approach outlined in Eriksson et al. 2013 (51). The scalar active stress in each cell was computed using the active contraction model by Land et al. 2017, 2018 (52, 53). Default active stress parameters were employed, apart from maximal active tension (*T*_*ref*_), which was set at 280 kPa to maintain a physiological ejection fraction of approximately 30% for all considered patient cases. To incorporate the impact of fibrosis on active tension generation, we multiplied the maximal active tension by a fixed factor of 0.5, to account for a reduced cardiac contraction in fibrotic tissue based on limited experimental data (49, 54). **Figure 1B** illustrates the prominent impact on contraction (as measured by physical deformation and strain development) when the biomechanical changes associated with hypocontractility, increased stiffness, or both are incorporated in a simple tissue geometry.

After achieving steady-state conditions using the 0D single-cell model to generate the initial conditions for the 3D model, we conducted a 3D volumetric EM simulation for one beat lasting 1,000 ms. After completing the EM simulations, we extracted the endocardial surfaces from the geometries along with their deformation over time. These extracted surfaces were subsequently used for the CFD simulations described below.

#### 2.2.2. Computational fluid dynamics model

The blood flow resulting from the simulated motion of patient-specific anatomies and fibrotic maps was modeled using the in-house code TUCAN (55). This code discretizes the incompressible Navier-Stokes equations in space using centered, second-order, finite differences. It integrates them in time using a fractional step method embedded within a low-storage, three-stage, semi-implicit Runge–Kutta scheme. It uses a cubic Eulerian (i.e., fixed) computational domain 13 × 13 × 13 cm^3^ and a staggered Cartesian mesh of size Δ*x*_*N*_ = 0.51 *mm*, resulting in LA computational fluid domains with 16.8 million mesh cells. The time stepping resolution dt was set to keep the Courant-Friedrichs-Lewy number below *CFL*_*max*_ = 0.28. Blood was modeled as a Newtonian fluid with constant kinematic viscosity, *ν* = 0.04 *cm*^2^*/s*.

The flow was coupled to the motion of the LA walls using an immersed boundary method (56). This method produces localized volumetric forces in the Eulerian mesh to match the LA wall velocity at the time-dependent positions of the endocardium obtained from the biomechanics simulations, which is discretized in a Lagrangian mesh (blue points in **Figure 1C**). The flow quantities are interpolated between the Eulerian and Lagrangian meshes using a regularized delta function kernel introduced by Peskin (57). A similar approach is used to impose inflow boundary conditions (BCs) at each pulmonary vein (PV) using the red points in **Figure 1C**, as previously described (58-60). The total PV inflow rate is obtained from mass conservation in the LA considering the change of volume in time of LA and the flow leaving the LA through the mitral valve annulus (i.e., outflow or left ventricle volume change in time). This total flow rate is split between the four PVs based on the ratio between their respective area and the total area of the PVs to prescribe the same velocity through each PV. LA outflow is simulated by not enforcing boundary conditions at the Lagrangian points used to model the mitral valve (MV) at timepoints where the valve is open (pink region of **Figure 1C**). Free-slip BCs are imposed at the outer edges of the Eulerian cubic mesh.

To simulate blood flow, we ran five cycles using a coarse spatial resolution (Δ*x*_*C*_ ≈ 0.9 *mm*). Then, the flow variables are interpolated into a fluid mesh with nominal resolution Δ*x*_*N*_ and ran five more cycles. This approach reduced the computational burden of the simulations by calculating the initial flow transient with the coarse spatial resolution and converging the velocity field in nominal resolution. The results presented were time-averaged for the last two cycles (from t/T = 8 to 10) once the cycle-to-cycle mean velocity variation is smaller than 6.5%.

#### 2.2.3. Coupling CFD and biomechanics models

The Lagrangian CFD mesh representing the endocardium was obtained from the biomechanics model as follows. An in-house code first generated a triangular surface mesh comprising the biomechanics model’s endocardial nodes at rest. This mesh was then refined to match the spatial resolution of the CFD code, and the endocardial displacements obtained by the biomechanics model are interpolated. The Lagrangian CFD mesh was completed by adding homogeneous planar triangulations at the mitral valve annulus (MVA) and pulmonary vein inlets. These operations were performed at 20 instants of the 60 bpm cardiac cycle separated by constant intervals of 0.05 s, representing “key time frames” of the cycle. This is temporal discretization is equal or finer than the method used for prior atrial flow simulations performed with TUCAN using Lagrangian meshes obtained from patient-specific 4D-CT images (58-60). Once the Lagrangian meshes representing key time frames of the cardiac cycle were obtained, the subsequent steps were identical to those used in previous TUCAN studies (58-60). In short, we registered the meshes using a coherent point drift algorithm (61) and apply Fourier interpolation in time to match the CFD temporal resolution.

### 2.3. Simulation of biomechanical fibrosis effects on LA booster function

The normal LA performs three mechanical functions. While the mitral valve is closed and the atrial myocardium relaxed, the LA acts as a passive reservoir, storing kinetic energy of blood returning from the lungs in the form of myocardial elastic energy. This energy is transferred back onto the flow when the LA recoils and the mitral valve is open, with the LA acting as a conduit for pulmonary vein blood to flow directly into the left ventricle. When the atrial myocardium contracts, the LA acts as a booster pump, actively delivering blood into the left ventricle. (62). Atrial fibrosis affects these three functions in an interdependent manner (i.e., it typically lowers reservoir and booster function and raises conduit function) by reducing the myocardium’s contractility and increasing its stiffness (16). We have previously shown that LA hemodynamics can be sensitive to the relative importance of reservoir, conduit, and booster functions (58), making it difficult to disentangle how fibrosis affects hemodynamics without separating its effects on each function. Consequently, and to illustrate how multi-physics models can be used to test mechanistic hypotheses, we focused on booster function, which can be modulated by both the active and passive myocardial properties.

At the beginning of each simulation, the LA contracted rapidly against a constant pressure of 10 mmHg and the LA emptied through the MVA (Baseline flow in **Figure 4A-B** upper rows, t/T < 0.2 - t < 0.15). This phase was followed by a slower relaxation phase (Baseline flow in Figure 4A-B lower rows, 0.2 < t/T < 0.51 - 0.15 < t/T < 0.46), during which the mitral valve was closed and the LA filled with blood coming from the pulmonary veins, passively recovering its initial volume. The duration of this cycle was set to be T = 1 s.

To dissect the biomechanical effects of fibrosis, we ran four different states by independently modifying stiffness and active tension in the fibrotic regions of each patient-specific anatomical model. We considered a state in which passive myocardial stiffness was increased five-fold (48) but activation tension was kept normal in fibrotic regions (*stiff-fibrotic*), another where myocardial stiffness was normal but activation tension generation was decreased two-fold (*hypocontractile-fibrotic*), and a state where both effects were included (*fibrotic*). For comparison, we also considered a *non-fibrotic* baseline state that rescued normal values of both biomechanical parameters throughout the whole LA. In addition, we ran simulations in the high-fibrosis models where both fibrosis effects were included, but fibrotic regions were digitally removed either in the LA body or the LAA (**Figure 7**). These simulations aimed to investigate how atrial flow is affected by the spatial distribution of fibrosis and the interactions between the flow in the LA and the LAA cavities. Considering all possible combinations of fibrosis distributions and parameter specifications, we ran a total of 20 multi-physics simulations in four patient-specific models.

### 2.4. Comparative analysis of hemodynamic metrics on volumetric Lagrangian meshes

Point-wise comparison of regional electromechanical and biomechanical metrics across different specifications of the same anatomical model is straightforward, since these variables are all discretized on a common Lagrangian mesh where each node corresponds to a material point in the myocardium. However, a similar comparison for hemodynamic metrics, which are discretized on a volumetric Cartesian Eulerian mesh that does not conform to the moving LA anatomy, is more involved. To address this challenge, we created patient-specific volumetric Lagrangian meshes by solving the biomechanical model inside the whole LA volume. This approach produces time-varying tetrahedral meshes with a constant number of elements registered across different instants of the cardiac cycle, conforming to each patient’s atrial time-dependent anatomy. To perform statistical comparisons of hemodynamic metrics on a node-by-node basis, the results from the CFD simulations were mapped into the volumetric Lagrangian meshes using the same 2_nd_-order interpolator used to discretize the Navier Stokes equations on the Eulerian mesh.

The flow kinetic energy, i.e., 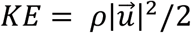 where *ρ* is the fluid’s density and 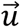 is the velocity vector, has been extensively used in previous studies to evaluate atrial hemodynamics (60, 63-65). To quantify the local hemodynamic changes induced by fibrosis, we normalized the *KE* at each point of the volumetric Lagrangian mesh and instant of time, i.e., 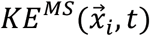 where the super index *MS* indicates a fibrotic mechanical state (*stiff-fibrotic, hypocontractile-fibrotic*, or *fibrotic*), with the value of the flow kinetic energy at the same Lagrangian point and instant of time in the *non-fibrotic* state. The normalized kinetic energy, defined as 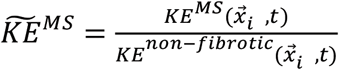 is lower than one in regions where fibrosis-induced effects lead to slower flow, and vice versa.

In addition to local, point-wise comparisons, we defined *KE*_*RoI*_ (*t*) to evaluate regional changes in flow *KE* by averaging in space over a region of interest in the volumetric Lagrangian mesh (ROI). Last, we normalized the *KE* with the fluid density, time-averaged ROI volume in the whole phase-averaged cycle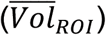, and heart rate (HR) to define a non-dimensional kinetic energy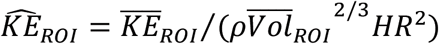. This metric facilitated comparison of flows in different patient-specific LA geometries. We chose as ROIs the atrial body, the LAA (the most frequent site of atrial thrombosis), and a blood pool region adjacent the endocardium (uniform 0.25 cm thickness) to assess near-endocardial vs. global hemodynamic effects of myocardial fibrosis.

## 3. Results

### 3.1. LA electrical activation from EP simulations

The differences in conductivity values between fibrotic and non-fibrotic tissue gave rise to modest changes in the tissue-scale electrical activity of the LA. In the sequence of maps of electrical activation shown in **Figure 2A**, the electrical stimulus is delivered in the region between the LAA and the left superior pulmonary vein, proximal to the area where Bachmann’s bundle would typically penetrate the LA myocardium. The presence of fibrosis caused subtle changes in activation sequence in both cases shown. The slowing effect arising from impaired transverse conductivity in fibrotic areas is more appreciable in the Fib41 subject compared to Fib07, but in both cases the overall effect is modest (excitation delays corresponding to <10% of the total activation time; reflected in histograms of activation time on right side of **Figure 2A**). In Fib41, conduction delays were impacted throughout the LA whereas the changes were localized to the largest patch of fibrotic tissue in the Fib07 case.

**Figure 2.**
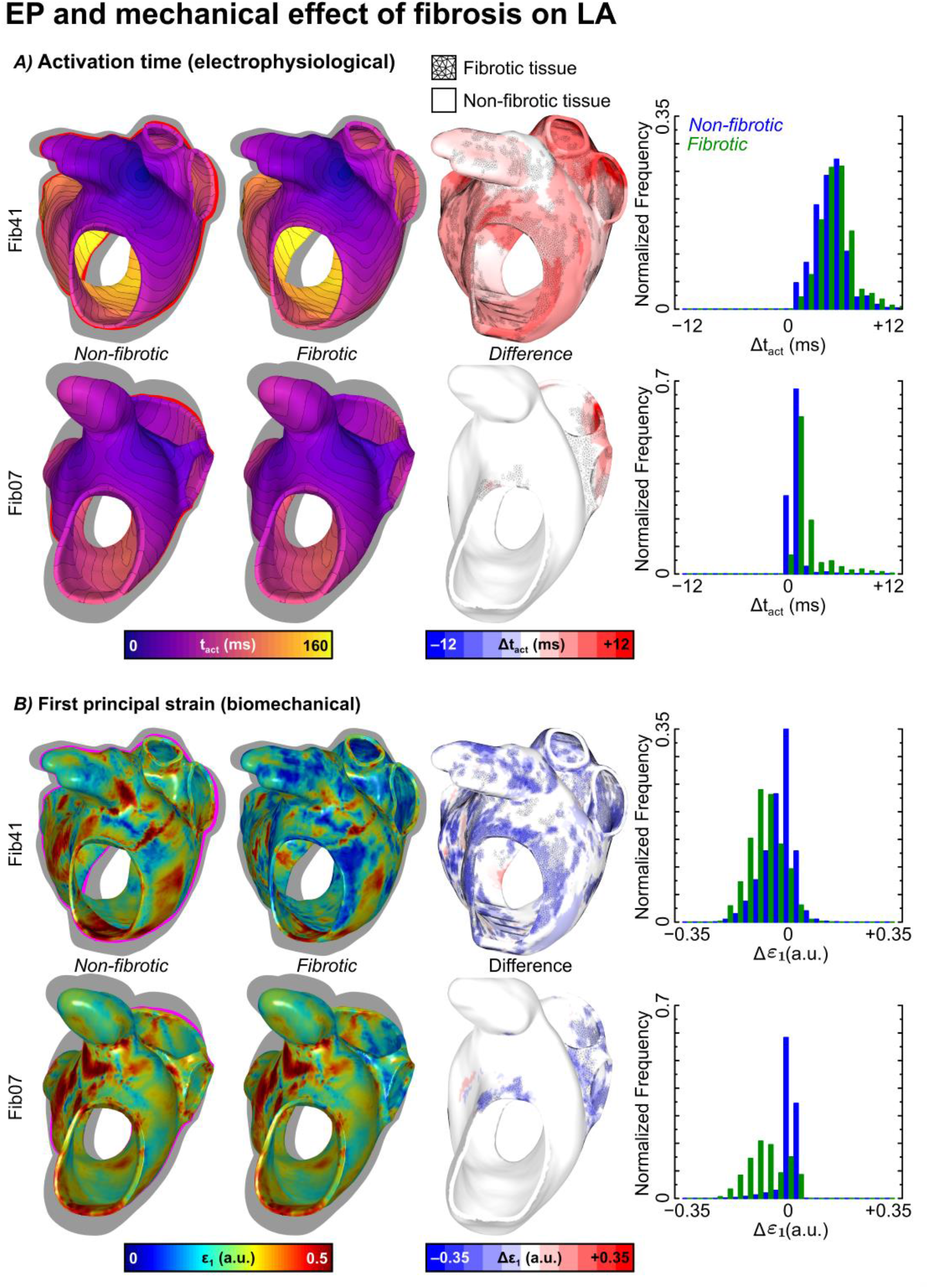
EP and mechanical effect of fibrosis on LA. **A)** Electrical activation time snapshots at maximum atrial contraction of subjects Fib41 (upper row) and Fib07 (lower row). At each row, the same view is shown for non-fibrotic simulations with equal conductivity values in fibrotic and non-fibrotic tissue (1^st^ column) and simulations with altered conductivity values in fibrotic tissue (2^nd^ column). Differences in activation time, *t*_*act*_, (column 3^rd^) are included in the same view for visual qualitative comparison, and histograms of those differences (4^th^ column) in fibrotic and non-fibrotic tissue for qualitative comparison. Fibrotic regions are identified with tessellated black areas on the myocardium maps showing activation time differences. **B)** First principal strain, *ε*_1_, snapshots on myocardium surface at maximum atrial contraction of subjects Fib41 (upper row) and Fib07 (lower row). At each row, the same view is shown for *non-fibrotic* (1^st^ column) and *fibrotic* (2^nd^ column) states. Differences in strain between states are shown as differences in activation time in **A)**. To ease the visualization of differences in LA contraction between states, the myocardium position at maximum atrial contraction of the *fibrotic* state is shown in pink color below the strain snapshots of the *non-fibrotic* state. Tessellated black areas also identify fibrotic regions on the myocardium maps showing strain differences.

### 3.2. Mechanical derangement due to fibrosis

Subtle changes in myocardial electrical propagation in EP simulations alter activation times for the biomechanics model, introducing a slight delay. The various states of the mechanical model that we tested (i.e., altering cardiomyocyte contraction and tissue stiffness) modified the organ-scale mechanical response much more dramatically. **Figure 2B** illustrates how these alterations affected the spatiotemporal pattern of myocardial deformation at peak LA contraction by mapping the first (highest) principal strain (*ε*_1_) on the same high-fibrosis (Fib41) and low-fibrosis (Fib07) patient-specific models of **Figure 2A**. The mechanical effects of fibrosis in our models were more pronounced in high-fibrosis patients compared to those with low fibrosis burdens, aligning with expectations. Specifically, subject Fib41 (**Figure 2B**, upper row) had significantly lower ε_1_ values throughout the LA for the *fibrotic* compared to *non-fibrotic* configurations. In the *non-fibrotic* configuration, a larger fraction of the LA surface is in the range 0.25–0.5 compared to the *fibrotic* configuration. The global differences in ε_1_ between the *fibrotic* and *non-fibrotic* configurations for subject Fib07 (lower row) were minor, but there were noteworthy local differences near the left PVs in the vicinity of fibrotic tissue.

The spatial LA contraction over the cardiac cycle can be qualitatively appreciated by comparing the gray silhouettes representing undeformed model shapes with the colored maps at maximum LA contraction in **Figure 2B**. LA contraction differences between mechanical states are depicted using magenta outlines of the LA endocardium of *fibrotic state* in *non-fibrotic* colored maps. For instance, in the case of subject Fib07 (lower row), the magenta regions highlight the posterior and septal walls of the LA as areas in which impaired contraction resulted in deformation differences between the *fibrotic* and *non-fibrotic* configurations. In contrast, for subject Fib41 in which fibrotic remodeling was much more pervasive throughout the chamber, the magenta outline is visible around all regions of the silhouette, indicating a high level of global impairment in the chamber’s ability to contrast in the *fibrotic* compared to the *non-fibrotic* configuration. These differences are borne out in our quantitative analysis via ε_1_ histograms (right panels of **Figure 2B**) for fibrotic and non-fibrotic myocardial regions. For both subjects considered here, the histograms associated with ε_1_ values in fibrotic areas were left-shifted (i.e., weaker local strain values) compared to those for non-fibrotic regions. Comparing these histograms with those in **Figure 2A** highlights the fact that the biomechanical consequences of fibrosis in these models were much more striking than the electrophysiological effects.

Detailed schematics for all four fibrotic LA models are shown in **Figure 3A**, including annotations of fibrosis burden in the LA body and LAA. Differences in electromechanics discussed in **Figure 2** corresponded to the temporal profiles of LA and LAA chamber volumes, presented for all simulations in **Figure 3B**. The minimum LA and LAA volumes were obtained in the *non-fibrotic* configuration for all patient-specific models, consistent with these atria contracting more compared to those in the *fibrotic* configuration. To quantify how fibrosis affected overall contractility, we calculated each model’s chamber emptying fraction (EF), a variable used clinically to quantify global mechanical function of the LA, versus fibrosis burden. Moreover, to analyze the independent and synergistic effects of fibrotic stiffening and hypocontractility, we normalized EFs in the *stiff-fibrotic, hypocontractile-fibrotic*, and *fibrotic* configurations with the baseline, *non-fibrotic* EF. These normalized values 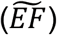 are plotted in **Figures 3C-D**. Mechanically impaired atrial function, associated with low values of EF, resulted in 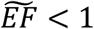 for all fibrotic simulations both in the LA body and the LAA. Furthermore, 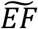 decreased monotonically with % fibrosis burden. Comparing the *stiff-fibrotic* and *hypocontractile-fibrotic* configurations, we found that hypocontractility had a weaker effect on 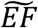 than stiffness in all cases except the LA body of subject Fib41.

**Figure 3.**
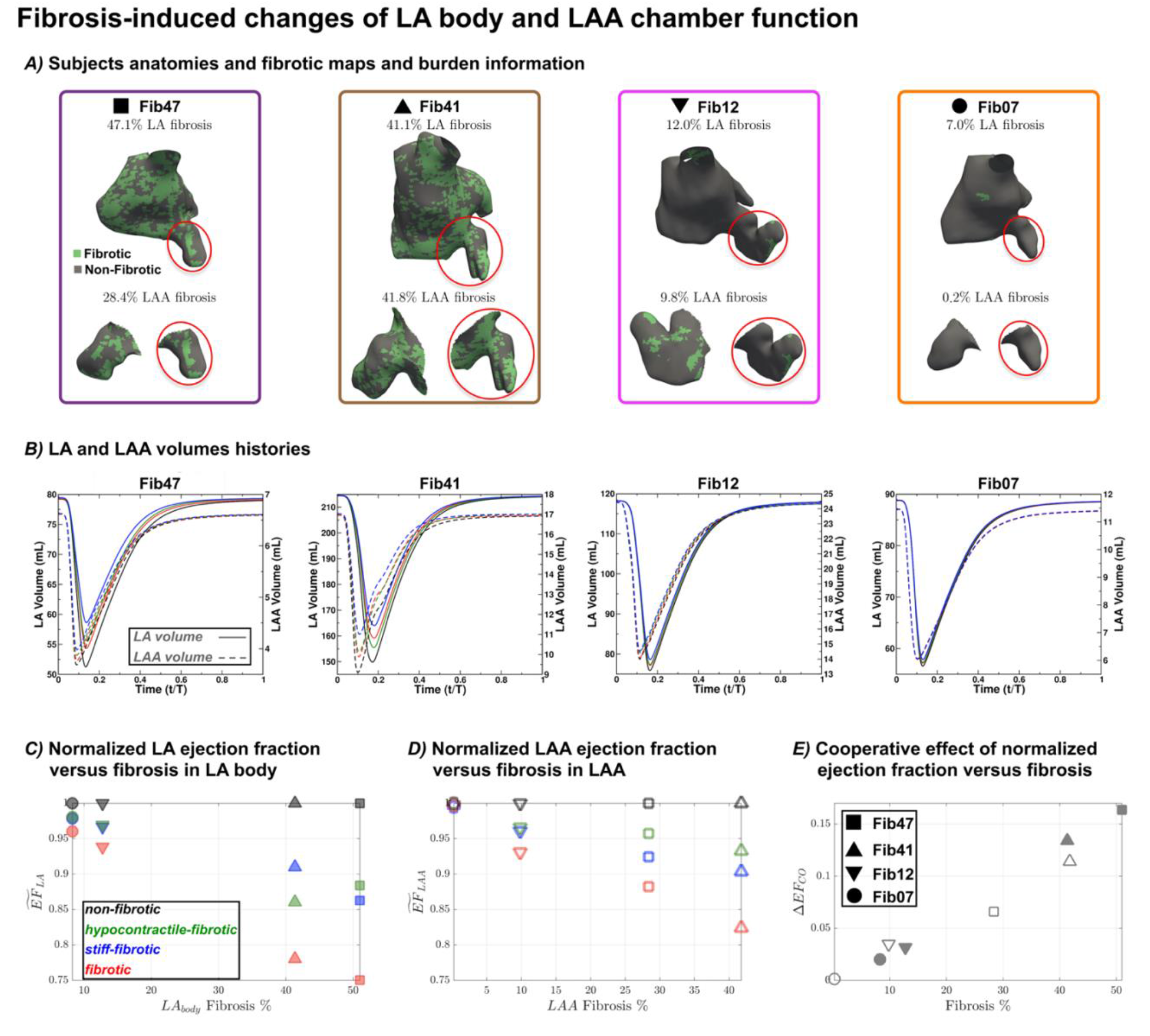
Fibrosis-induced changes of LA body and LAA chamber function. **A)** Patient-specific LA anatomy models, including fibrotic maps (fibrotic tissue in green and non-fibrotic in gray) and LA and LAA fibrosis %. **B)** Time histories of LA (solid line) and LAA (dashed line) volumes for all patient-specific models. LA and LAA volumes of the four mechanical states are represented in black (*non-fibrotic*), green (*hypocontractile-fibrotic*), blue (*stiff-fibrotic*), and red (*fibrotic*). Patient-specific models are ordered from left to right by percentage of fibrosis burden in the LA. **C)** LA body emptying fraction normalized with its non-fibrotic state value for each patient-specific model, 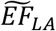, plotted vs. each model’s % fibrosis burden in the LA body. The blue, green, and red symbols represent the *stiff-fibrotic, hypocontractile-fibrotic*, and *fibrotic* states, respectively. **D)** Same as panel A for the LAA. **E)** Coefficient of fibrotic mechanical cooperativity, Δ*EF*_*co*_, on the EF of the LA body (solid symbols) and the LAA (hollow symbols) as a function of each model’s % fibrosis burden. See main text for definition of Δ*EF*_*co*_. Subjects are identified with markers (▪→Fib47, ▴→Fib41, ▾→ Fib12, •→ Fib07).

**Figure 4.**
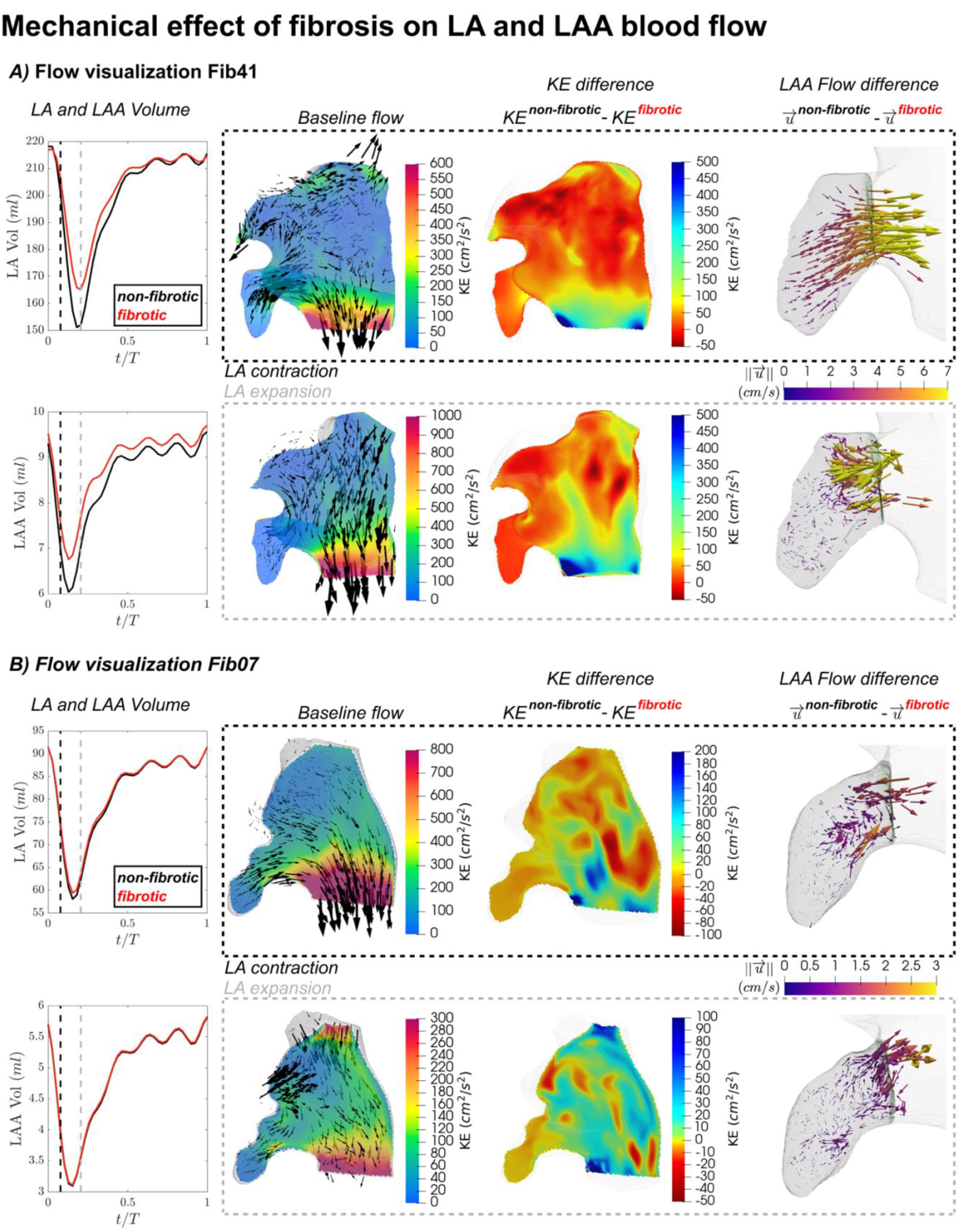
Mechanical effect of fibrosis on LA and LAA blood flow. **A)** Case with high fibrosis burden (Fib41). **B)** Case with low fibrosis burden (Fib07). Each panel displays the time histories of LA (upper row) and LAA (lower row) volumes for *fibrotic* (red) and *non-fibrotic* (black) mechanical states on the left column. For each subject, the blood flow visualizations are shown at two time instants: peak LAA contraction rate (upper row, black rectangle) and peak LAA expansion rate (lower row, gray rectangle). These time instants are indicated with black and gray vertical dashed lines in the LA and LAA volume plots on the left-hand side of each panel. On the left side of each rectangle, an instant snapshot of the kinetic energy (*KE*) of the *non-fibrotic* state is shown in a plane section dissecting LA body and LAA. Back arrows scaled with the velocity magnitude are included in the snapshot to visualize the flow direction (arrows in lower row of panel B are 1.5X larger than in upper row). On the center of each rectangle, the difference in kinetic energy between *non-fibrotic* and *fibrotic* states is represented in the same plane section used for *KE* snapshots. On the right-hand side of each rectangle, an LAA view with vectors showing the velocity difference between *non-fibrotic* and *fibrotic* states is presented. Arrows in this LAA view are scaled and colored with the velocity difference magnitude.

Of note, 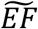 in the *fibrotic* configuration decreased with fibrosis burden, amplifying the reduction observed in the *stiff-fibrotic* and *hypocontractile-fibrotic* configurations in a non-linear fashion. This prompted us to calculate a coefficient of fibrotic mechanical cooperativity, defined as:

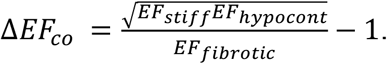

A positive value of this metric indicates that fibrotic stiffness and hypocontractility amplify each other, since the *fibrotic* configuration exhibits lower EF values than the geometric mean of the *stiff-fibrotic* and *hypocontractile-fibrotic* configurations; conversely, Δ*EF*_*co*_ < 0 implies mutual attenuation between the two types of biomechanical derangement. This cooperativity coefficient is plotted vs. % fibrosis burden for all four models in **Figure 3E**, demonstrating that the two fibrosis effects amplified each other more intensely as the fibrosis burden increased.

### 3.3. Effect of fibrosis on atrial flow patterns

Changes in stiffness and contractility of fibrotic myocardial modified LA endocardium motion, ultimately affecting LA blood flow patterns. Flow visualization and point-wise comparison of flow metrics using the volumetric Lagrangian mesh defined in section 2.4 provided insight into these hemodynamic alterations. **Figure 4** displays blood flow velocity maps and flow *KE* at two instants of the simulations corresponding to peak contraction and expansion rate of the LAA (indicated by the black and grey vertical dashed lines in the volume vs. time curves of **Figure 4A-B**). These time points correspond roughly to the early contraction and expansion of the LA. The same two models presented in **Figure 2** (Fib41 and Fib07) are shown in **Figure 4**.

Subject Fib41’s *non-fibrotic* baseline flow at peak LAA contraction rate displayed a straight jet emanating from the contracting LAA that merged with the strong mitral outflow jet. Concomitantly, the incipient contraction of the LA body caused backflow through the PVs (**Figure 4A**, top row). These flow patterns are commonly observed in CFD simulations driven by time-resolved atrial segmentations obtained from clinical 4D-CT (58-60, 66, 67). Adding the mechanical effects of fibrosis significantly disturbed LA flow patterns in this high-fibrosis subject. The flow *KE* for the *fibrotic* configuration decreased throughout the LA, most appreciably in the aforementioned regions where baseline flow was strong (see 3_rd_ column of **Figure 4A**). The 3D vectorial representation of the velocity differences in the LAA(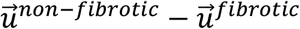, fourth column of **Figure 4A**) shows that the LAA emptying jet was weakened by ∼5 cm/s in the *fibrotic* state. At peak LAA expansion rate (**Figure 4A**, bottom row), the Fib41 model’s baseline flow pattern displayed a weaker MVA outflow jet, while blood flowing through the PV replenished the LA and flowed from the LAA ostium to the distal LAA. Like the previous time point, adding fibrosis effects decreased the flow *KE* most significantly in these regions and, to a lesser extent, throughout the entire LA. Of note, 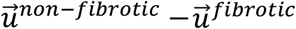 showed a coherent inflow pattern at the LAA ostium, also of magnitude ∼5 cm/s, suggesting that LAA filling flow was slower in the *fibrotic* state than in the *non-fibrotic* state.

In the Fib07 subject (**Figure 4B**), the overall patterns of the *non-fibrotic* baseline flow were comparable with those seen in the subject Fib41’s baseline flow. On the other hand, consistent with the relatively modest mechanical derangement caused by fibrosis, the flow *KE* decreased less significantly in this subject than in Fib41. However, there were appreciable *KE* differences between the *fibrotic* and *non-fibrotic* configurations across the entire atrium, even if most of subject Fib07’s fibrotic tissue was localized near the left PVs. In particular, flow inside the LAA suffered noticeable alterations, albeit 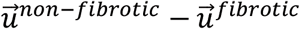 reached lower magnitudes (∼1 cm/s) and did not exhibit coherent patterns during LAA emptying or filling compared to the high-fibrosis case.

We quantified how fibrosis disturbed LA hemodynamics by computing the flow *KE* inside the LA and LAA for all fibrotic configurations in all models, and normalized each value with the corresponding value of the *non-fibrotic* configuration (see 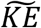 definition in section 2.4). To evaluate whether fibrotic effects on blood flow were global or confined to the vicinity of the endocardium, we performed similar analyses for the entire LA chamber and a region within 0.25 cm of the LA endocardium. Time-averaged 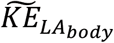 and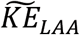 values are plotted in **Figure 5** and for completeness time-averaged 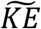 distributions are shown in **Figure SI 1-2**.

**Figure 5.**
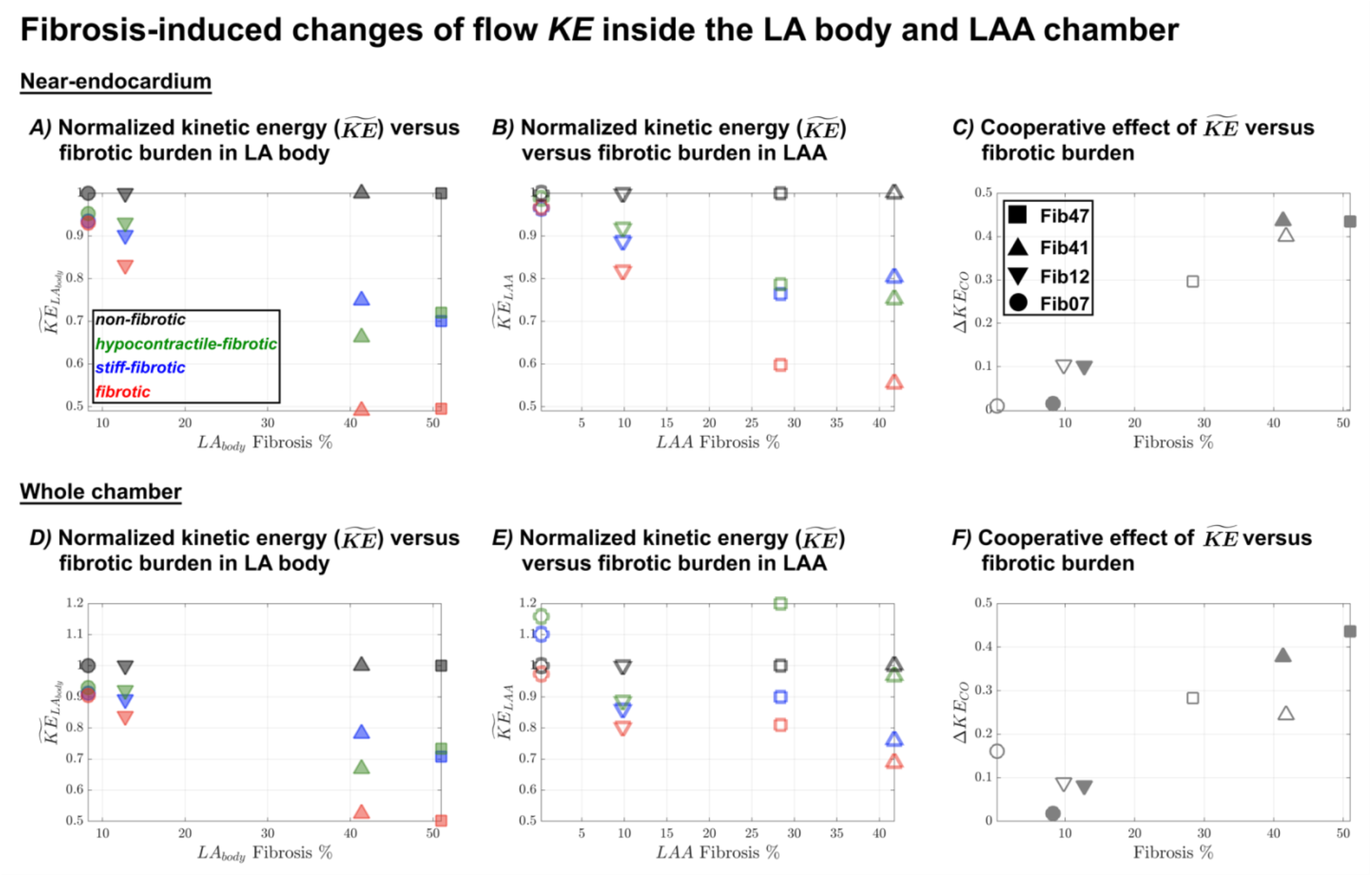
Fibrosis-induced changes of flow *KE* inside the LA body and LAA chamber. Near-endocardium results are shown in panels **A-C** and whole chamber results in panels **D-F. A)** and **D)** Total flow kinetic energy in the LA body normalized with its *non-fibrotic* state value for each patient-specific model, 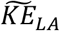, plotted vs. each model’s % fibrosis burden in the LA body. The black, blue, green, and red symbols represent the *non-fibrotic, stiff-fibrotic, hypocontractile-fibrotic*, and *fibrotic* states, respectively. **B)** and **E)** Same as panel **A** and **C** for the LAA. **C)** and **F)** Coefficient of fibrotic hemodynamic cooperativity, Δ*KE*_*co*_, on the *KE* of the LA body (solid symbols) and the LAA (hollow symbols) as a function of each model’s % fibrosis burden. See main text for definition of Δ*KE*_*co*_. Subjects are identified with markers (▪→Fib47, ▴→Fib41, ▾→ Fib12, •→ Fib07). Values of 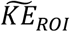 presented in this figure are time-averaged in the whole phase-averaged cycle and spatially averaged in the *RoI* of the surface Lagrangian mesh for near-endocardium results and the volumetric Lagrangian mesh for whole chamber results.

Consistent with flow velocities matching tissue velocities at the endocardial border (i.e., the no-slip boundary condition), the dependence of near-endocardial *KE* on fibrosis burden was very similar to that of the LA EF reported above. The normalized near-endocardial flow kinetic energy 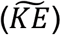 decreased with increasing fibrosis burden in both the LA body (**Figure 5A**) and LAA (**Figure 5B**); these changes were prominent, with 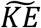 values falling approximately two-fold in the patient-specific models with high fibrosis. Significant cooperativity, as quantified by the coefficient

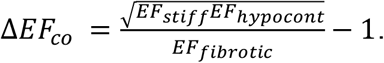

was also observed (**Figure 5C**). When we considered flow *KE* in the whole volume of the LA body (**Figure 5D**), we found a very similar trend, suggesting the fibrotic derangement of myocardial mechanics globally affects LA hemodynamics. On the other hand, the whole-LAA flow *KE* responded differently to fibrosis (**Figure 5E**). First, there were a few instances with fibrotic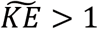, implying that flow inside the LAA became stronger on average in a few simulations with stiffness-only or hypocontractility-only configurations, but this phenomenon was never observed for the more pathophysiologically relevant *fibrotic* configuration. Second, 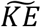 was less sensitive to fibrosis burden than the in the LA body. Third, Δ*KE*_*co*_ was lower for the LAA than for the LA body in all models and did not vary significantly with fibrosis burden (**Figure 5F**), suggesting that effects of stiffness and hypocontractility amplified each other in a more complex, multi-factorial manner inside the LAA.

### 3.5. Fibrotic effects on atrial hemodynamics can be reduced to changes in global chamber function for the atrial body but not for the atrial appendage

Finally, we explored whether changes in flow *KE* caused by fibrosis-induced loss of mechanical function could be indexed to global LA chamber function or whether additional parameters, like chamber geometry, fibrosis distribution, etc., were needed to reproduce these changes. **Figure 6A** shows the non-dimensional mean flow *KE* in the LA body vs. this chamber’s emptying fraction, i.e.,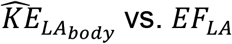. Consistent with the results shown in previous figures, 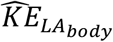 increased monotonically with *EF*_*LA*_, and both quantities decreased in configurations with fibrosis effects for all patient-specific models. A striking feature of this plot is that all the data points collapsed tightly around the line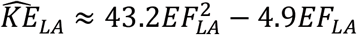, suggesting that the biomechanical effects of fibrosis on the hemodynamics inside the LA body were fully captured by the changes in this chamber’s global function. When we analyzed the same variables in the LAA (**Figure 6B**), we observed a similar trend, but the data were more scattered, suggesting that the hemodynamics of the LAA are more sensitive to additional factors like LAA morphology, the spatial distribution of fibrosis, or the interactions between the flow in the LA and the LAA cavities.

**Figure 6.**
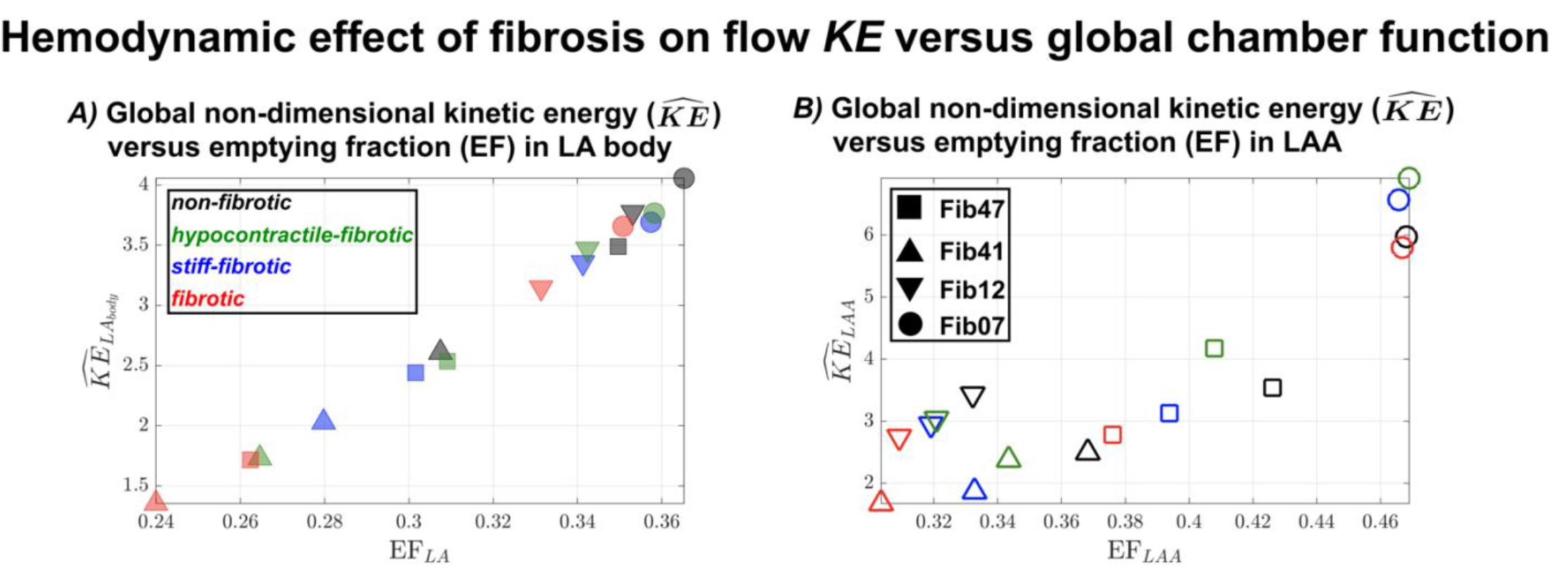
Hemodynamic effect of fibrosis on flow *KE* versus global chamber function. Non-dimensional flow kinetic energy in the LA body,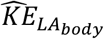, vs. LA emptying fraction. The black, blue, green, and red symbols represent the *non-fibrotic, stiff-fibrotic, hypocontractile-fibrotic*, and *fibrotic* states, respectively. Subjects are identified with markers (▪→Fib47, ▴→Fib41, ▾→ Fib12, •→ Fib07). **B)** Same as panel **A** for the LAA. Values of 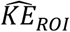are time-averaged in the whole phase-averaged cycle and spatially averaged in the *RoI* (see definition in section 2.4).

**Figure 7.**
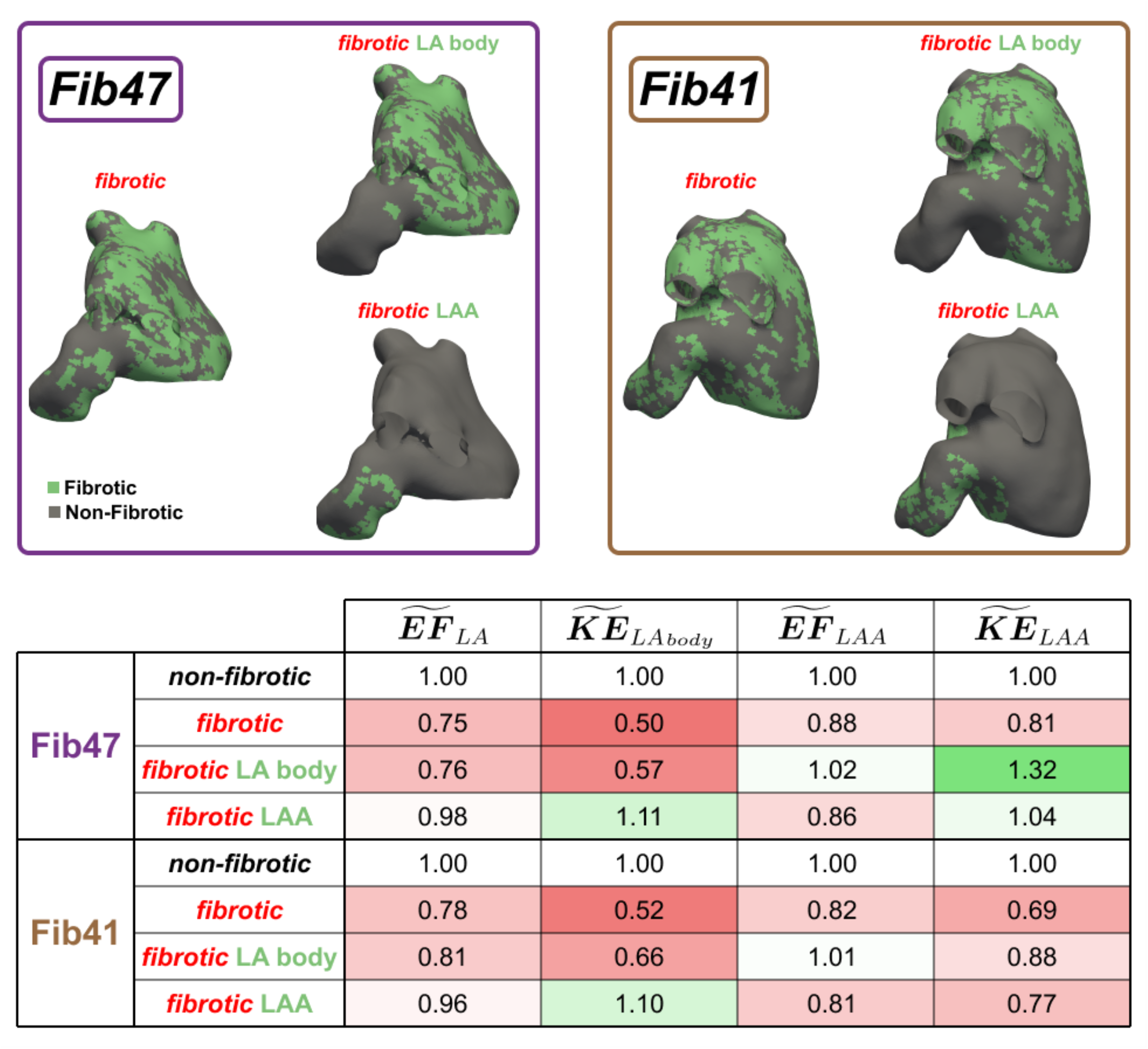
Effect of fibrosis on global chamber function and flow *KE*. 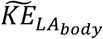 and 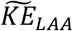 are the normalized flow kinetic energy (see definition in section 2.4) in the left atrial (LA) body and the left atrial appendage (LAA), respectively. 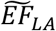 and 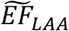 are the left atrium and LAA normalized emptying fraction, respectively. Values below (i.e., < 1) and above (i.e., > 1) *non-fibrotic* state’s reference value are colored in red and green, respectively. Coloring fading is used to indicate proximity to the reference value. Values of 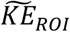 presented in this table are time-averaged in the whole phase-averaged cycle and spatially averaged in the *RoI* of the volumetric Lagrangian mesh.

To further investigate this idea, we conducted simulations in which mechanical changes were induced in fibrotic points of LA body or LAA independently. **Figure 7** displays the fibrotic maps of these simulations. To maximize the impact of fibrosis-induced mechanical changes on the flow, we only simulated the *fibrotic* state since this state exhibited larger *KE* differences with respect to *non-fibrotic* simulations. Similarly, we only considered the cases with higher fibrotic burdens (Fib41 and Fib47). The differences in global mechanic function and flow dynamics are summarized in **Figure 7**. Of note, 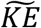 in the LAA is strongly affected when fibrosis is considered exclusively in the LA body, despite the small change in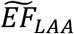. The subject-dependent behavior of 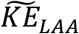 (increasing in subject Fib47 while decreasing for Fib41) highlights the intricate interaction between global LA body mechanical function and flow with LAA flow. Also, and as expected, LAA fibrosis alone also impacted 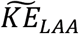 due to a reduced LAA contraction (i.e., lower values of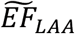). However, the increase of 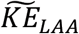 in subject Fib47 suggests that the effects produced by LA and LAA body fibrosis in LAA flow can be comparable.

## 4. Discussion

This work presents a modular multi-physics, multi-scale, computational framework to investigate how fibrotic changes in myocardial electrophysiology and biomechanics impact blood flow inside the left atrium. This framework constitutes a weakly coupled electromechanical model that interfaces with a 3D computational fluid dynamics (CFD) module. At each instant of the cardiac cycle, the CFD solver uses an immersed boundary method to ensure that blood velocity on the endocardial surfaces matches tissue velocity. We used this computational framework with patient-specific data from magnetic resonance imaging (MRI) as input, including fibrotic tissue maps from late gadolinium enhancement (LGE). In fibrotic tissue regions, we used modified values for parameters describing the myocardium’s electrical conduction, stiffness, and contractile tension. Conversely, normal parameter values were used in non-fibrotic regions. To our knowledge, this is the first computational framework to explicitly model myocardial fibrosis and resolve its effects on LA hemodynamics. The framework allows for modifying parameters controlling a single mechanism in fibrotic regions while keeping non-fibrotic values for all other parameters. We leveraged this capability to investigate how fibrotic changes in tissue stiffness and activation tension, both independently and together, affect flow inside the left atrium. To establish closely matched controls for the fibrotic patient-specific models, baseline simulations were performed on identical anatomies with fibrosis turned off in the models. Moreover, to investigate how total fibrosis burden, defined as the percentage of tissue with fibrosis throughout the LA, affects blood flow, we considered four patient-specific cases with low (7%, 12%) and high (41%, 47%) burdens of fibrosis.

The link between atrial fibrosis and stroke is supported by ample clinical and experimental evidence (10, 13, 14, 68-70). However, the exact mechanisms by which fibrosis leads to an increased risk of stroke are far from understood. Fibrotic remodeling modifies LA electrophysiology and mechanical function through processes occurring at different spatial scales (15, 16). One prominent effect in myocardium afflicted with AF-associated fibrosis is impaired conduction, particularly in the transverse direction (71). Normal cardiac tissue can be characterized as an anisotropic material with a stiffer passive response and active tension generation along the direction of myocardial fibers (72). In fibrotic regions, the active tension produced by cardiomyocyte contraction decreases due to a reduction of force-producing myocytes (73). Furthermore, increased stiffness is due to excessive collagen deposition (74, 75). Both factors contribute to decreased tissue contractility in all directions when balancing active and passive tension. Reduced contractility could significantly affect blood flow patterns, creating stagnant regions associated with high thrombogenesis risk (14).

The approach used in this study to represent the bioelectric and biomechanical consequences of fibrosis was based on the combination of patient-specific imaging data with generic parameterization changes derived from previously reported physiological properties derived from experimental testing of cardiac tissue. This approach binarizes the changes attributed to fibrosis since each finite element in our representation of the LA is assigned either fibrotic or non-fibrotic electromechanical properties. As evidenced by the large number of prior studies that have used this approach (76-78), it is a necessary simplification since model parameterizations representing areas with different degrees of local fibrotic remodeling have yet to be available. Notably, this does not mean fibrotic and non-fibrotic regions in our simulations behave monolithically – due to electrotonic interaction between adjacent regions enforced by the bioelectric propagation formulation, functional gradients in electrophysiological properties emerge dynamically as steady-state conditions are approached. Similarly, we observed high variations in mechanical strain within both fibrotic and non-fibrotic tissue.

Turning fibrosis on in the models significantly decreased myocardial strain along the cardiac cycle, especially at peak contraction, in agreement with the clinical evidence of an inverse correlation between fibrosis burden and atrial strain (79, 80). The impact of fibrosis on electrical propagation was relatively modest since we opted not to include cell-scale remodeling of ion channel expression levels in these regions; this allowed us to analyze the purely biomechanical consequences of fibrosis. In atria with low fibrosis burdens, remodeling-related strain effects were mostly confined to fibrotic regions, while in highly fibrotic atria, the effects were also seen in non-fibrotic regions. In our *in-silico* models, these non-local effects are necessarily caused by the balance of forces within the tissue, suggesting that mechanics could play a significant role in propagating fibrotic impairment of myocardial strain. *In vivo*, additional processes such as fibrosis-mediated inflammation (81), likely exacerbated by epicardial fat (82, 83), can propagate the fibrotic derangement of myocardial contraction.

When integrated over the whole chamber, the fibrotic impairment of myocardial strain reduced the active LA emptying fraction. The reduction was more significant as overall fibrosis burden increased, reaching a 20-25% drop in EF for fibrosis burdens of 40-50% (81, 84). These results are in good agreement with clinical findings that elderly subjects with high LA LGE area coverage (≥17 cm_2_) have a 20% lower active LA EF than subjects with low LA LGE coverage (<17 cm_2_) (85). While either increasing stiffness or decreasing active tension in fibrotic patches independently reduced LA EF, including the two fibrotic derangements together in the models lowered the LA EF even further. We found this cooperative effect to be more pronounced when fibrosis burden was increased, a behavior likely due to the non-linear behavior of the myocardial stiffness.

The effect of fibrosis on blood flow through the left atrium and its appendage has been scarcely addressed in the literature. Left atrial fibrotic mapping can be obtained on a patient-specific basis using LGE-MRI (86, 87), and there are several imaging modalities to assess LAA filling and emptying flows (88-91). However, few studies assess both fibrosis and flow (92), and even fewer examine how fibrosis is associated with flow patterns or markers of blood stasis, like spontaneous echocardiographic contrast (87). To our knowledge, the only previous CFD investigation on atrial hemodynamics in fibrotic atria is the study by Paliwal et al. (32). They reported increased values of hemodynamic metrics associated with endothelial injury and slow flow near fibrotic regions on the endocardium in subjects with a higher fibrosis burden, consistent with our observations. However, Paliwal et al.’s (32) simulations used a simplified endocardial motion model based on each patient’s global LA function instead of resolving how fibrotic maps influence regional myocardial biomechanics. This modeling choice might have obscured the non-local effects of fibrosis on LA and LAA hemodynamics found in our study.

Our CFD analysis of fibrotic models showed that fibrosis significantly altered atrial flow patterns throughout the whole atrial chamber and, particularly, inside the LAA. Flow disturbances were most significant in regions of fast flow such as the mitral outflow and the PV inlet regions and, for the most part, consisted of flow deceleration. Remarkably, the filling and emptying velocities at the LAA ostium weakened significantly in highly fibrotic cases, implying that LAA clearance is impaired and LAA stasis increases in fibrotic atria. This result agrees with pulsed Doppler measurements obtained in the LAA of a canine model of atrial fibrosis (93). It also agrees with the increased rate of spontaneous echocardiographic contrast inside the LAA of highly fibrotic atria (87) and is consistent with the widely reported clinical association between atrial fibrosis and ischemic stroke risk (10). Near the endocardium, the fibrosis-induced changes in flow velocity mirrored the regional changes in LA wall velocity via the no-slip boundary condition imposed in the simulations. Concordantly, we found the near-endocardial flow *KE* to vary with fibrosis burden very much like chamber emptying fraction. However, fibrotic alterations of atrial hemodynamics were not confined to the vicinity of the endocardium. Significant non-local effects of boundary conditions are to be expected since blood must satisfy mass conservation and momentum balance, and atrial flow has significant inertia as indicated by its Reynolds number hovering over the thousands.

In the LA body, fibrotic effects in hemodynamics should be relatively uninvolved, as the loss of this chamber’s flow *KE* was explained uniquely by the fibrosis-induced changes in its emptying fraction, regardless of atrial anatomy or spatial distribution of fibrosis. This observation agrees with previous findings that atrial flow inside the LA body is largely determined by the chamber’s global volume balance, so that hemodynamic metrics like its average residence time are well approximated by simple laws involving chamber volume and stroke volume, independent of PV inflow split or blood rheological properties (59, 60). The relatively simple morphology and global hemodynamics of the LA body offers some justification for Paliwal et al.’s (32) simplified models for endocardial motion.

However, our multi-physics models suggest that fibrotic flow disturbances are multi-factorial in the LAA, the most common site of atrial thrombosis. Consequently, the risk of LAA thrombosis has been shown to depend on additional factors like LA and LAA morphology (94-96), and it is reasonable to expect this risk to also depend on the spatial distribution of fibrosis rather than on its global effect on LAA emptying fraction. Being an intricately shaped tubular sac misaligned with the overall direction of atrial flow, LAA hemodynamics can be highly sensitive to anatomical parameters and boundary conditions, including endocardial motion in distant regions of the LA. Our simulations suggest that, despite the general trend for LAA flow kinetic energy to decrease with fibrosis burden, this variation is less robust than in the LA body, even showing increases of LAA flow kinetic energy on the *hypocontractile-fibrotic* and *stiff-fibrotic* states simulations in Fib07 and the *hypocontractile-fibrotic* state simulation in Fib47 (see **Figure 5**). This is not unreasonable, as one can easily imagine several scenarios where LAA flow could become faster in stiffer or hypocontractile atria. For instance, if the proximal LAA stiffens more than the distal LAA, flow velocities through the ostium could increase to accommodate a similar volume flux through a smaller orifice. Likewise, reduced LAA contractility could curb the cyclic manifestation of geometrical features, such as high LAA bending angles, which are associated with slow LAA flow (97). Furthermore, our results from simulations considering fibrosis in LA body or LAA separately suggest a tangled interaction between the flow of both cavities. LAA blood stasis is influenced by the dynamics of the jets produced during LAA contraction and expansion and by the recirculating flow from the LA body. LAA jets are subject to mass conservation inside this blind-ended cavity, while flow from the LA body interacts with LAA flow through shearing and fluid exchange processes near the ostium. Further work is needed to continue characterizing the dependence of LAA hemodynamics on fibrosis and its interplay with additional parameters like LAA morphology.

### Study Limitations and Future Work

The number of patient-specific models considered in this study was small (N=4), even if we ran four different simulations per patient-specific case to generate the *non-fibrotic* baseline state as well as the *hypocontractile-fibrotic, stiff-fibrotic*, and *fibrotic* states. To mitigate this issue, we selected two models with modest fibrosis burdens (7% and 12%) and two models with severe fibrosis burdens (41% and 47%). This approach allowed for detecting differences between low-fibrosis and high-fibrosis models at the expense of neglecting intermediate fibrosis burdens.

In addition to the pumping function created by active expansion and contraction of the atrial myocardium, the atrium operates passively as a reservoir and a conduit. When both pulmonary vein pressure and LV pressure exceed LA pressure, the mitral valve is closed, and the blood incoming to the LA causes this chamber to expand passively, acting as a *reservoir*. When LA pressure rises above LV pressure, the mitral valve opens, and the blood stored in the LA can flow to the LV as the LA recoils passively. At the same time, the LA serves as a *conduit*, allowing venous pulmonary return to flow directly into the LV. Clinical evidence suggests that fibrosis impairs left atrial strain associated with these three functions (84, 85). However, in a first effort towards understanding the physical phenomena linking fibrosis and aberrant hemodynamics in the LA, we focused on active atrial pumping. Thus, we modeled LV pressure as constant throughout the cardiac cycle and set the maximum LA volume to be the same in the fibrotic and non-fibrotic simulations for each patient-specific model. Future simulations should address more realistic atrial cycles including reservoir and conduit functions.

Many different approaches to representing the EP consequences of fibrosis have been explored (77). There is no consensus on which single method is most suitable, but all approaches tend to produce similar organ-scale consequences (e.g., reentrant driver localization). Our EP model accounts for fibrosis by decreasing the conduction velocity (CV) in fibrotic tissue, but no differences in cell-scale action potential are considered between fibrotic and non-fibrotic regions. As discussed in Methods, this minimalist approach to modeling the EP of fibrotic tissue was a deliberate design choice in the present study since we sought to assess the specific organ-scale biomechanical consequences of deranged stiffness and active tension generation in fibrotic tissue. Incorporating further changes to the EP ionic model (e.g., reduced L-type Ca_2+_ current) would have significantly complicated the clean analysis of these relationships. Given the uncertainty surrounding whether these changes are needed to reproduce biophysically plausible behavior, we believe this decision is justified.

Multiplying the active tension and passive stiffness by fixed factors at all fibrotic points of all subjects’ anatomical models is a substantial simplification. Myocardial fibrosis occurs in several different forms (17). For instance, in response to irregular mechanical stresses caused by AFib or pressure overload, the matrix around cardiomyocytes expands, leading to “diffuse fibrosis.” On the other hand, when cardiomyocytes die due to, e.g., hypoxia or catheter ablation, a new extracellular matrix is created in their place, leading to “replacement fibrosis.” In each fibrosis class, collagenous matrix and cardiomyocytes adopt different spatial patterns likely to have varying impacts on local contractility and stiffness. However, there is limited experimental data on this potential relationship.

In our multi-physics computational framework, the electromechanical and CFD models are one-way coupled, i.e., spatial pressure fluctuations created by the flow do not deform the myocardium. This lack of two-way fluid-structure interaction could be particularly concerning in the left atrium, which sustains lower mean chamber pressures than the left ventricle. Nonetheless, the flow patterns predicted by our simulations compare well with those obtained when our same CFD solver is fed by time-resolved atrial models obtained from 4D computed tomography scans (58-60). As far as we know, there are no previous simulations resolving how fibrosis influences myocardial biomechanics and LA hemodynamics. This paucity justifies the present study as a first effort to understand this influence.

Blood is primarily composed of plasma and erythrocytes. While plasma behaves as a Newtonian fluid of constant viscosity, erythrocytes tend to aggregate in regions of slow flow, conferring blood a hematocrit-dependent shear-thinning non-Newtonian rheology. Previous studies including our own have shown that non-Newtonian rheology could significantly affect the flow inside the LAA, where blood sustains low shear rates for extended periods of time (98). However, we considered blood as a Newtonian fluid to reduce the parametric space required to set the simulations and focus on the effect of mechanical properties changes in fibrotic tissue.

## 5. Conclusion

Multi-physics, multi-scale simulations coupling electrophysiology, biomechanics, and blood flow were performed for the first time on patient-specific left atrial models including fibrotic maps from LGE-MRI. In these simulations, fibrotic and, to a lesser degree, non-fibrotic tissue experienced impaired motion, leading to decreased LA and LAA emptying fractions. Consequently, atrial flow patterns were disturbed throughout the atrial chamber, weakening the LAA filling and emptying jets. These disturbances were noticeable even in models with low fibrosis burden, becoming more severe as the extent of disease-related remodeling increased. This work illustrates how multi-physics models leveraging patient-specific anatomy and fibrotic distribution from non-invasive medical imaging can contribute to understanding the mechanisms by which fibrosis affects atrial function and promotes thrombosis.

## Supporting information

Figure SI 1 and Figure SI 2 used to dissect the hemodynamic effects of fibrotic tissue stiffening and hypocontractility

## Data availability statement

The original contributions presented in the study are included in Results and Supplementary Information. Patient-specific left atrial anatomical models and fibrotic maps are publicly available at Dryad (https://doi.org/10.5061/dryad.d7wm37q9h). Simulation data requires over 1TB of storage per subject, making it unsuitable for repository sharing. Data will be shared upon reasonable request for non-commercial use. Any further inquiries can be directed to the corresponding author.

## References

1. Feigin VL, Brainin M, Norrving B, Martins S, Sacco RL, Hacke W, et al. World Stroke Organization (WSO): Global Stroke Fact Sheet 2022. International Journal of Stroke. 2022;17(1):18–29. doi: 10.1177/17474930211065917.

2. Waolff P. Atrial fibrillation as an independent risk factor for stroke. Stroke. 1991;22:983–8.

3. Lippi G, Sanchis-Gomar F, Cervellin G. Global epidemiology of atrial fibrillation: an increasing epidemic and public health challenge. International Journal of Stroke. 2021;16(2):217–21.

4. Lamassa M, Di Carlo A, Pracucci G, Basile AM, Trefoloni G, Vanni P, et al. Characteristics, outcome, and care of stroke associated with atrial fibrillation in Europe: data from a multicenter multinational hospital–based registry (The European Community Stroke Project). Stroke. 2001;32(2):392–8.

5. Lin H-J, Wolf PA, Kelly-Hayes M, Beiser AS, Kase CS, Benjamin EJ, et al. Stroke severity in atrial fibrillation: the Framingham Study. Stroke. 1996;27(10):1760–4.

6. Crowther MA, Warkentin TE. Bleeding risk and the management of bleeding complications in patients undergoing anticoagulant therapy: focus on new anticoagulant agents. Blood, The Journal of the American Society of Hematology. 2008;111(10):4871–9.

7. Sun Y, Ling Y, Chen Z, Wang Z, Li T, Tong Q, et al. Finding low CHA2DS2-VASc scores unreliable? Why not give morphological and hemodynamic methods a try? Frontiers in Cardiovascular Medicine. 2023;9:1032736.

8. Wynn T. Cellular and molecular mechanisms of fibrosis. The Journal of Pathology: A Journal of the Pathological Society of Great Britain and Ireland. 2008;214(2):199–210.

9. Cunha PS, Laranjo S, Heijman J, Oliveira MM. The Atrium in Atrial Fibrillation–A Clinical Review on How to Manage Atrial Fibrotic Substrates. Frontiers in cardiovascular medicine. 2022;9:879984.

10. Daccarett M, Badger TJ, Akoum N, Burgon NS, Mahnkopf C, Vergara G, et al. Association of left atrial fibrosis detected by delayed-enhancement magnetic resonance imaging and the risk of stroke in patients with atrial fibrillation. Journal of the American College of Cardiology. 2011;57(7):831–8.

11. Gramley F, Lorenzen J, Knackstedt C, Rana OR, Saygili E, Frechen D, et al. Age-related atrial fibrosis. Age. 2009;31:27–38.

12. Akoum N, Mahnkopf C, Kholmovski EG, Brachmann J, Marrouche NF. Age and sex differences in atrial fibrosis among patients with atrial fibrillation. EP Europace. 2018;20(7):1086–92.

13. Tandon K, Tirschwell D, Longstreth W, Smith B, Akoum N. Embolic stroke of undetermined source correlates to atrial fibrosis without atrial fibrillation. Neurology. 2019;93(4):e381–e7.

14. Boyle PM, Del Álamo JC, Akoum N. Fibrosis, atrial fibrillation and stroke: clinical updates and emerging mechanistic models. Heart. 2021;107(2):99–105.

15. Frangogiannis NG. Cardiac fibrosis. Cardiovascular research. 2021;117(6):1450–88.

16. Kong P, Christia P, Frangogiannis NG. The pathogenesis of cardiac fibrosis. Cellular and molecular life sciences. 2014;71:549–74.

17. Telle Å, Bargellini C, Chahine Y, Del Álamo JC, Akoum N, Boyle PM. Personalized biomechanical insights in atrial fibrillation: opportunities & challenges. Expert Review of Cardiovascular Therapy. 2023;21(11):817–37. doi: 10.1080/14779072.2023.2273896.

18. Kirchhof P, Breithardt G, Aliot E, Al Khatib S, Apostolakis S, Auricchio A, et al. Personalized management of atrial fibrillation: Proceedings from the fourth Atrial Fibrillation competence NETwork/European Heart Rhythm Association consensus conference. Europace. 2013;15(11):1540–56.

19. Nordsletten D, Niederer S, Nash M, Hunter P, Smith N. Coupling multi-physics models to cardiac mechanics. Progress in biophysics and molecular biology. 2011;104(1-3):77–88.

20. Verzicco R. Electro-fluid-mechanics of the heart. Journal of Fluid Mechanics. 2022;941:P1.

21. Watanabe H, Sugiura S, Kafuku H, Hisada T. Multiphysics simulation of left ventricular filling dynamics using fluid-structure interaction finite element method. Biophysical journal. 2004;87(3):2074–85.

22. Sugiura S, Okada J-I, Washio T, Hisada T. UT-heart: A finite element model designed for the multiscale and multiphysics integration of our knowledge on the human heart. Computational Systems Biology in Medicine and Biotechnology: Methods and Protocols: Springer; 2022. p. 221–45.

23. Vigmond EJ, Clements C, McQueen DM, Peskin CS. Effect of bundle branch block on cardiac output: a whole heart simulation study. Progress in biophysics and molecular biology. 2008;97(2-3):520–42.

24. Quarteroni A, Lassila T, Rossi S, Ruiz-Baier R. Integrated heart—coupling multiscale and multiphysics models for the simulation of the cardiac function. Computer Methods in Applied Mechanics and Engineering. 2017;314:345–407.

25. Bucelli M, Zingaro A, Africa PC, Fumagalli I, Dede’ L, Quarteroni A. A mathematical model that integrates cardiac electrophysiology, mechanics, and fluid dynamics: Application to the human left heart. International Journal for Numerical Methods in Biomedical Engineering. 2023;39(3):e3678.

26. Zingaro A, Vergara C, Dede’ L, Regazzoni F, Quarteroni A. A comprehensive mathematical model for cardiac perfusion. Scientific Reports. 2023;13(1):14220.

27. Feng L, Gao H, Griffith B, Niederer S, Luo X. Analysis of a coupled fluid-structure interaction model of the left atrium and mitral valve. International journal for numerical methods in biomedical engineering. 2019;35(11):e3254.

28. Viola F, Meschini V, Verzicco R. Fluid–structure-electrophysiology interaction (FSEI) in the left-heart: a multi-way coupled computational model. European Journal of Mechanics-B/Fluids. 2020;79:212–32.

29. Choi YJ, Constantino J, Vedula V, Trayanova N, Mittal R. A new MRI-based model of heart function with coupled hemodynamics and application to normal and diseased canine left ventricles. Frontiers in bioengineering and biotechnology. 2015;3:140.

30. Viola F, Spandan V, Meschini V, Romero J, Fatica M, de Tullio MD, et al. FSEI-GPU: GPU accelerated simulations of the fluid–structure–electrophysiology interaction in the left heart. Computer physics communications. 2022;273:108248.

31. Viola F, Del Corso G, De Paulis R, Verzicco R. GPU accelerated digital twins of the human heart open new routes for cardiovascular research. Scientific Reports. 2023;13(1):8230.

32. Paliwal N, Ali RL, Salvador M, O’Hara R, Yu R, Daimee UA, et al. Presence of left atrial fibrosis may contribute to aberrant hemodynamics and increased risk of stroke in atrial fibrillation patients. Frontiers in Physiology. 2021:684.

33. Bifulco SF, Scott GD, Sarairah S, Birjandian Z, Roney CH, Niederer SA, et al. Computational modeling identifies embolic stroke of undetermined source patients with potential arrhythmic substrate. Elife. 2021;10:e64213.

34. Bifulco SF, Macheret F, Scott GD, Akoum N, Boyle PM. Explainable machine learning to predict anchored reentry substrate created by persistent atrial fibrillation ablation in computational models. Journal of the American Heart Association. 2023;12(16):e030500.

35. Macheret F, Bifulco SF, Scott GD, Kwan KT, Chahine Y, Afroze T, et al. Comparing Inducibility of Re-Entrant Arrhythmia in Patient-Specific Computational Models to Clinical Atrial Fibrillation Phenotypes. Clinical Electrophysiology. 2023;9(10):2149–62.

36. Boyle PM, Sarairah S, Kwan KT, Scott GD, Mohamedali F, Anderson CA, et al. Elevated fibrosis burden as assessed by MRI predicts cryoballoon ablation failure. Journal of Cardiovascular Electrophysiology. 2023;34(2):302–12.

37. Marrouche NF, Wazni O, McGann C, Greene T, Dean JM, Dagher L, et al. Effect of MRI-guided fibrosis ablation vs conventional catheter ablation on atrial arrhythmia recurrence in patients with persistent atrial fibrillation: the DECAAF II randomized clinical trial. Jama. 2022;327(23):2296–305.

38. Marrouche NF, Wilber D, Hindricks G, Jais P, Akoum N, Marchlinski F, et al. Association of atrial tissue fibrosis identified by delayed enhancement MRI and atrial fibrillation catheter ablation: the DECAAF study. Jama. 2014;311(5):498–506.

39. Neic A, Gsell MA, Karabelas E, Prassl AJ, Plank G. Automating image-based mesh generation and manipulation tasks in cardiac modeling workflows using meshtool. SoftwareX. 2020;11:100454.

40. Roney CH, Bendikas R, Pashakhanloo F, Corrado C, Vigmond EJ, McVeigh ER, et al. Constructing a human atrial fibre atlas. Annals of biomedical engineering. 2021;49:233–50.

41. Augustin CM, Neic A, Liebmann M, Prassl AJ, Niederer SA, Haase G, et al. Anatomically accurate high resolution modeling of human whole heart electromechanics: a strongly scalable algebraic multigrid solver method for nonlinear deformation. Journal of computational physics. 2016;305:622–46.

42. Plank G, Loewe A, Neic A, Augustin C, Huang Y-L, Gsell MA, et al. The openCARP simulation environment for cardiac electrophysiology. Computer Methods and Programs in Biomedicine. 2021;208:106223.

43. Courtemanche M, Ramirez RJ, Nattel S. Ionic mechanisms underlying human atrial action potential properties: insights from a mathematical model. American Journal of Physiology-Heart and Circulatory Physiology. 1998;275(1):H301–H21.

44. Krummen DE, Bayer JD, Ho J, Ho G, Smetak MR, Clopton P, et al. Mechanisms of human atrial fibrillation initiation: clinical and computational studies of repolarization restitution and activation latency. Circulation: Arrhythmia and Electrophysiology. 2012;5(6):1149–59.

45. Boyle PM, Zahid S, Trayanova NA. Towards personalized computational modelling of the fibrotic substrate for atrial arrhythmia. EP Europace. 2016;18(uppl_4):iv136–iv45.

46. Potse M, Dubé B, Richer J, Vinet A, Gulrajani RM. A comparison of monodomain and bidomain reaction-diffusion models for action potential propagation in the human heart. IEEE Transactions on Biomedical Engineering. 2006;53(12):2425–35.

47. Augustin CM, Fastl TE, Neic A, Bellini C, Whitaker J, Rajani R, et al. The impact of wall thickness and curvature on wall stress in patient-specific electromechanical models of the left atrium. Biomechanics and modeling in mechanobiology. 2020;19(3):1015–34.

48. Chaturvedi RR, Herron T, Simmons R, Shore D, Kumar P, Sethia B, et al. Passive stiffness of myocardium from congenital heart disease and implications for diastole. Circulation. 2010;121(8):979–88.

49. Mastikhina O, Moon B-U, Williams K, Hatkar R, Gustafson D, Mourad O, et al. Human cardiac fibrosis-on-a-chip model recapitulates disease hallmarks and can serve as a platform for drug testing. Biomaterials. 2020;233:119741.

50. Marx L, Niestrawska JA, Gsell MA, Caforio F, Plank G, Augustin CM. Robust and efficient fixed-point algorithm for the inverse elastostatic problem to identify myocardial passive material parameters and the unloaded reference configuration. Journal of computational physics. 2022;463:111266.

51. Eriksson TS, Prassl AJ, Plank G, Holzapfel GA. Influence of myocardial fiber/sheet orientations on left ventricular mechanical contraction. Mathematics and Mechanics of Solids. 2013;18(6):592–606.

52. Land S, Park-Holohan S-J, Smith NP, Dos Remedios CG, Kentish JC, Niederer SA. A model of cardiac contraction based on novel measurements of tension development in human cardiomyocytes. Journal of molecular and cellular cardiology. 2017;106:68–83.

53. Land S, Niederer SA. Influence of atrial contraction dynamics on cardiac function. International journal for numerical methods in biomedical engineering. 2018;34(3):e2931.

54. Travers JG, Kamal FA, Robbins J, Yutzey KE, Blaxall BC. Cardiac fibrosis: the fibroblast awakens. Circulation research. 2016;118(6):1021–40.

55. Moriche M. A numerical study on the aerodynamic forces and the wake stability of flapping flight at low Reynolds number [PhD thesis]: Universidad Carlos III de Madrid; 2017.

56. Uhlmann M. An immersed boundary method with direct forcing for the simulation of particulate flows. Journal of computational physics. 2005;209(2):448–76.

57. Peskin CS. The immersed boundary method. Acta numerica. 2002;11:479–517.

58. García-Villalba M, Rossini L, Gonzalo A, Vigneault D, Martinez-Legazpi P, Durán E, et al. Demonstration of patient-specific simulations to assess left atrial appendage thrombogenesis risk. Frontiers in physiology. 2021;12:596596.

59. Gonzalo A, García-Villalba M, Rossini L, Durán E, Vigneault D, Martínez-Legazpi P, et al. Non-Newtonian blood rheology impacts left atrial stasis in patient-specific simulations. International Journal for Numerical Methods in Biomedical Engineering. 2022;38(6):e3597.

60. Durán E, García-Villalba M, Martínez-Legazpi P, Gonzalo A, McVeigh E, Kahn AM, et al. Pulmonary vein flow split effects in patient-specific simulations of left atrial flow. Computers in Biology and Medicine. 2023:107128.

61. Myronenko A, Song X. Point set registration: Coherent point drift. IEEE transactions on pattern analysis and machine intelligence. 2010;32(12):2262–75.

62. Blume GG, Mcleod CJ, Barnes ME, Seward JB, Pellikka PA, Bastiansen PM, et al. Left atrial function: physiology, assessment, and clinical implications. European Journal of Echocardiography. 2011;12(6):421–30.

63. Triposkiadis F, Harbas C, Sitafidis G, Skoularigis J, Demopoulos V, Kelepeshis G. Echocardiographic assessment of left atrial ejection force and kinetic energy in chronic heart failure. The international journal of cardiovascular imaging. 2008;24:15–22.

64. Arvidsson PM, Töger J, Heiberg E, Carlsson M, Arheden H. Quantification of left and right atrial kinetic energy using four-dimensional intracardiac magnetic resonance imaging flow measurements. Journal of applied physiology. 2013;114(10):1472–81.

65. Zingaro A, Ahmad Z, Kholmovski E, Sakata K, Dede’ L, Morris AK, et al. A comprehensive stroke risk assessment by combining atrial computational fluid dynamics simulations and functional patient data. bioRxiv. 2024:2024.01. 11.575156.

66. Vedula V, George R, Younes L, Mittal R. Hemodynamics in the left atrium and its effect on ventricular flow patterns. Journal of biomechanical engineering. 2015;137(11):111003.

67. Lantz J, Henriksson L, Persson A, Karlsson M, Ebbers T. Patient-specific simulation of cardiac blood flow from high-resolution computed tomography. Journal of biomechanical engineering. 2016;138(12):121004.

68. King JB, Azadani PN, Suksaranjit P, Bress AP, Witt DM, Han FT, et al. Left atrial fibrosis and risk of cerebrovascular and cardiovascular events in patients with atrial fibrillation. Journal of the American College of Cardiology. 2017;70(11):1311–21.

69. Kühnlein P, Mahnkopf C, Majersik JJ, Wilson BD, Mitlacher M, Tirschwell D, et al. Atrial fibrosis in embolic stroke of undetermined source: a multicenter study. European journal of neurology. 2021;28(11):3634–9.

70. Mahnkopf C, Kwon Y, Akoum N. Atrial fibrosis, ischaemic stroke and atrial fibrillation. Arrhythmia & Electrophysiology Review. 2021;10(4):225.

71. King JH, Huang CL-H, Fraser JA. Determinants of myocardial conduction velocity: implications for arrhythmogenesis. Frontiers in physiology. 2013;4:154.

72. Avazmohammadi R, Soares JS, Li DS, Raut SS, Gorman RC, Sacks MS. A contemporary look at biomechanical models of myocardium. Annual review of biomedical engineering. 2019;21:417–42.

73. Wang EY, Rafatian N, Zhao Y, Lee A, Lai BFL, Lu RX, et al. Biowire model of interstitial and focal cardiac fibrosis. ACS central science. 2019;5(7):1146–58.

74. Münch J, Abdelilah-Seyfried S. Sensing and responding of cardiomyocytes to changes of tissue stiffness in the diseased heart. Frontiers in Cell and Developmental Biology. 2021;9:642840.

75. Liu H, Fan P, Jin F, Huang G, Guo X, Xu F. Dynamic and static biomechanical traits of cardiac fibrosis. Frontiers in Bioengineering and Biotechnology. 2022;10:1042030.

76. Zahid S, Cochet H, Boyle PM, Schwarz EL, Whyte KN, Vigmond EJ, et al. Patient-derived models link re-entrant driver localization in atrial fibrillation to fibrosis spatial pattern. Cardiovascular research. 2016;110(3):443–54.

77. Roney CH, Bayer JD, Zahid S, Meo M, Boyle PM, Trayanova NA, et al. Modelling methodology of atrial fibrosis affects rotor dynamics and electrograms. EP Europace. 2016;18(uppl_4):iv146–iv55.

78. Boyle PM, Zghaib T, Zahid S, Ali RL, Deng D, Franceschi WH, et al. Computationally guided personalized targeted ablation of persistent atrial fibrillation. Nature biomedical engineering. 2019;3(11):870–9.

79. Kuppahally SS, Akoum N, Burgon NS, Badger TJ, Kholmovski EG, Vijayakumar S, et al. Left atrial strain and strain rate in patients with paroxysmal and persistent atrial fibrillation: relationship to left atrial structural remodeling detected by delayed-enhancement MRI. Circulation: Cardiovascular Imaging. 2010;3(3):231–9.

80. Lisi M, Mandoli GE, Cameli M, Pastore MC, Righini FM, Benfari G, et al. Left atrial strain by speckle tracking predicts atrial fibrosis in patients undergoing heart transplantation. European Heart Journal-Cardiovascular Imaging. 2022;23(6):829–35.

81. Thomas TP, Grisanti LA. The dynamic interplay between cardiac inflammation and fibrosis. Frontiers in physiology. 2020;11:529075.

82. Ng AC, Strudwick M, van der Geest RJ, Ng AC, Gillinder L, Goo SY, et al. Impact of epicardial adipose tissue, left ventricular myocardial fat content, and interstitial fibrosis on myocardial contractile function. Circulation: Cardiovascular Imaging. 2018;11(8):e007372.

83. Conte M, Petraglia L, Cabaro S, Valerio V, Poggio P, Pilato E, et al. Epicardial adipose tissue and cardiac arrhythmias: focus on atrial fibrillation. Frontiers in Cardiovascular Medicine. 2022;9:932262.

84. Hopman LH, Mulder MJ, van der Laan AM, Demirkiran A, Bhagirath P, van Rossum AC, et al. Impaired left atrial reservoir and conduit strain in patients with atrial fibrillation and extensive left atrial fibrosis. Journal of Cardiovascular Magnetic Resonance. 2021;23:1–10.

85. Olsen FJ, Bertelsen L, Vejlstrup N, Diederichsen SZ, Bjerregaard CL, Graff C, et al. Association between four-dimensional echocardiographic left atrial measures and left atrial fibrosis assessed by left atrial late gadolinium enhancement. European Heart Journal-Cardiovascular Imaging. 2023;24(1):152–61.

86. Doltra A, Stawowy P, Dietrich T, Schneeweis C, Fleck E, Kelle S. Magnetic resonance imaging of cardiovascular fibrosis and inflammation: from clinical practice to animal studies and back. BioMed research international. 2013;2013.

87. Akoum N, Fernandez G, Wilson B, Mcgann C, Kholmovski E, Marrouche N. Association of atrial fibrosis quantified using LGE-MRI with atrial appendage thrombus and spontaneous contrast on transesophageal echocardiography in patients with atrial fibrillation. Journal of cardiovascular electrophysiology. 2013;24(10):1104–9.

88. Romero J, Cao JJ, Garcia MJ, Taub CC. Cardiac imaging for assessment of left atrial appendage stasis and thrombosis. Nature Reviews Cardiology. 2014;11(8):470–80.

89. Fatkin D, Kelly RP, Feneley MP. Relations between left atrial appendage blood flow velocity, spontaneous echocardiographic contrast and thromboembolic risk in vivo. Journal of the American College of Cardiology. 1994;23(4):961–9.

90. Antonielli E, Pizzuti A, Pálinkás A, Tanga M, Gruber No, Michelassi C, et al. Clinical value of left atrial appendage flow for prediction of long-term sinus rhythm maintenance in patients with nonvalvular atrial fibrillation. Journal of the American College of Cardiology. 2002;39(9):1443–9.

91. Markl M, Lee DC, Furiasse N, Carr M, Foucar C, Ng J, et al. Left atrial and left atrial appendage 4D blood flow dynamics in atrial fibrillation. Circulation: Cardiovascular Imaging. 2016;9(9):e004984.

92. Hwang SH, Oh Y-W, Kim M-N, Park S-M, Shim WJ, Shim J, et al. Relationship between left atrial appendage emptying and left atrial function using cardiac magnetic resonance in patients with atrial fibrillation: comparison with transesophageal echocardiography. The International Journal of Cardiovascular Imaging. 2016;32:163–71.

93. Li Y, Li W, Yang B, Han W, Dong D, Xue J, et al. Effects of Cilazapril on atrial electrical, structural and functional remodeling in atrial fibrillation dogs. Journal of electrocardiology. 2007;40(1):100. e1-. e6.

94. Khurram IM, Dewire J, Mager M, Maqbool F, Zimmerman SL, Zipunnikov V, et al. Relationship between left atrial appendage morphology and stroke in patients with atrial fibrillation. Heart rhythm. 2013;10(12):1843–9.

95. Yamamoto M, Seo Y, Kawamatsu N, Sato K, Sugano A, Machino-Ohtsuka T, et al. Complex left atrial appendage morphology and left atrial appendage thrombus formation in patients with atrial fibrillation. Circulation: Cardiovascular Imaging. 2014;7(2):337–43.

96. Di Biase L, Natale A, Romero J. Thrombogenic and arrhythmogenic roles of the left atrial appendage in atrial fibrillation: clinical implications. Circulation. 2018;138(18):2036–50.

97. Yaghi S, Chang A, Ignacio G, Scher E, Panda N, Chu A, et al. Left atrial appendage morphology improves prediction of stagnant flow and stroke risk in atrial fibrillation. Circulation: Arrhythmia and Electrophysiology. 2020;13(2):e008074.

98. Sanatkhani S, Nedios S, Menon PG, Saba SF, Jain SK, Federspiel WJ, et al. Subject-specific factors affecting particle residence time distribution of left atrial appendage in atrial fibrillation: A computational model-based study. Frontiers in Cardiovascular Medicine. 2023;10:1070498.

