## Supplementary material for "Multi-physics simulations reveal hemodynamic impacts of patient-derived fibrosis-related changes in left atrial tissue mechanics": Figure SI 1 and Figure SI 2 used to dissect the hemodynamic effects of fibrotic tissue stiffening and hypocontractility

### Supplementary Information

#### S.I. 1. Dissecting the hemodynamic effects of fibrotic tissue stiffening and hypocontractility

To analyze the mechanical effects of fibrosis considered in our models, i.e., stiffening and hypocontractility, both independently and together, we determined the statistical distributions of flow $KE$ in the *stiff-fibrotic*, *hypocontractile-fibrotic*, and *fibrotic* states, normalized with the baseline *non-fibrotic* $KE$ at each point of the volumetric Lagrangian grid. **Figure SI 1** presents the probability density functions (p.d.f.s) of normalized $KE$, $p(\widetilde{KE})$, when pooling data from the LA body and the LAA. Furthermore, to evaluate whether mechanical effects of fibrosis are global or confined to the endocardial vicinity, we also computed $p(\widetilde{KE})$ pooling data from a 0.25 cm - thick near endocardial region. The two same patient-specific models considered in the previous figures (Fib41 and Fib07) were represented in **Figure SI 1**.

In subject Fib41, the near-endocardium and whole-volume p.d.f.s of $\widetilde{KE}$ were similar to each other in the LA body (left-hand side of **Figure SI 1A**), suggesting that fibrosis effects were not restricted to the endocardial region. Overall, the two p.d.f.s corresponding to the *stiff-fibrotic* and *hypocontractile-fibrotic* states were shifted to $\widetilde{KE} < 1$ values, indicating that both kinds of mechanical disruptions independently weakened the flow inside the LA. Furthemore, the shift was significantly more pronounced for the *fibrotic* state, with modal values of $\widetilde{KE}$ around 0.6, i.e., a 40% decrease. This trend was also observed in the Fib47 case (**Figure SI 2**), supporting that increased stiffness and reduced contractility of fibrotic tissue cooperatively disturb LA hemodynamics. In the LAA of the Fib41 model (right-hand side of **Figure SI 1A**), the volumetric and near-endocardial distributions of $\widetilde{KE}$ were also similar to each other except for the *hypocontractile-fibrotic* state. Like in the LA body, a cooperative effect between stiffness and hypocontractility was observed.

It could be argued that the pronounced global fibrotic effects observed in subject Fib41 should be expected, considering this subject's high fibrosis burden. In contrast, subject Fib07 (**Figure SI 1B**) provided an interesting opportunity to probe local vs. global effects because this subject's fibrotic tissue was mostly confined to a small region surrounding the left PVs. Specifically, subject Fib07's LAA was non-fibrotic, yet the LAA flow volumetric $\widetilde{KE}$ distributions decreased in the *stiff-fibrotic* and *fibrotic* states. Since subject Fib07's total fibrosis burden was low, the p.d.f.s of $\widetilde{KE}$ were shifted modestly peaking around $\widetilde{KE} = 0.9$.

Finally, a noteworthy difference between subject Fib07 and Fib41 was that the effects of increasing stiffness and reducing contractility in the Fib07 model did not seem to be cooperative. In the LA body, the three p.d.f.s corresponding to the *stiff-fibrotic, hypocontractile-fibrotic*, and *fibrotic* states were shifted towards the left by the same amount. In the LAA, the p.d.f. corresponding to the *fibrotic* and *stiff-fibrotic* states were shifted to the left by a similar amount, whereas the *hypocontractile-fibrotic* state was barely shifted.

### Dissecting the hemodynamic effects of fibrotic tissue stiffening and hypocontractility

##### *A)* Probability distribution function of normalized kinetic energy ($\widetilde{KE}$) Fib41

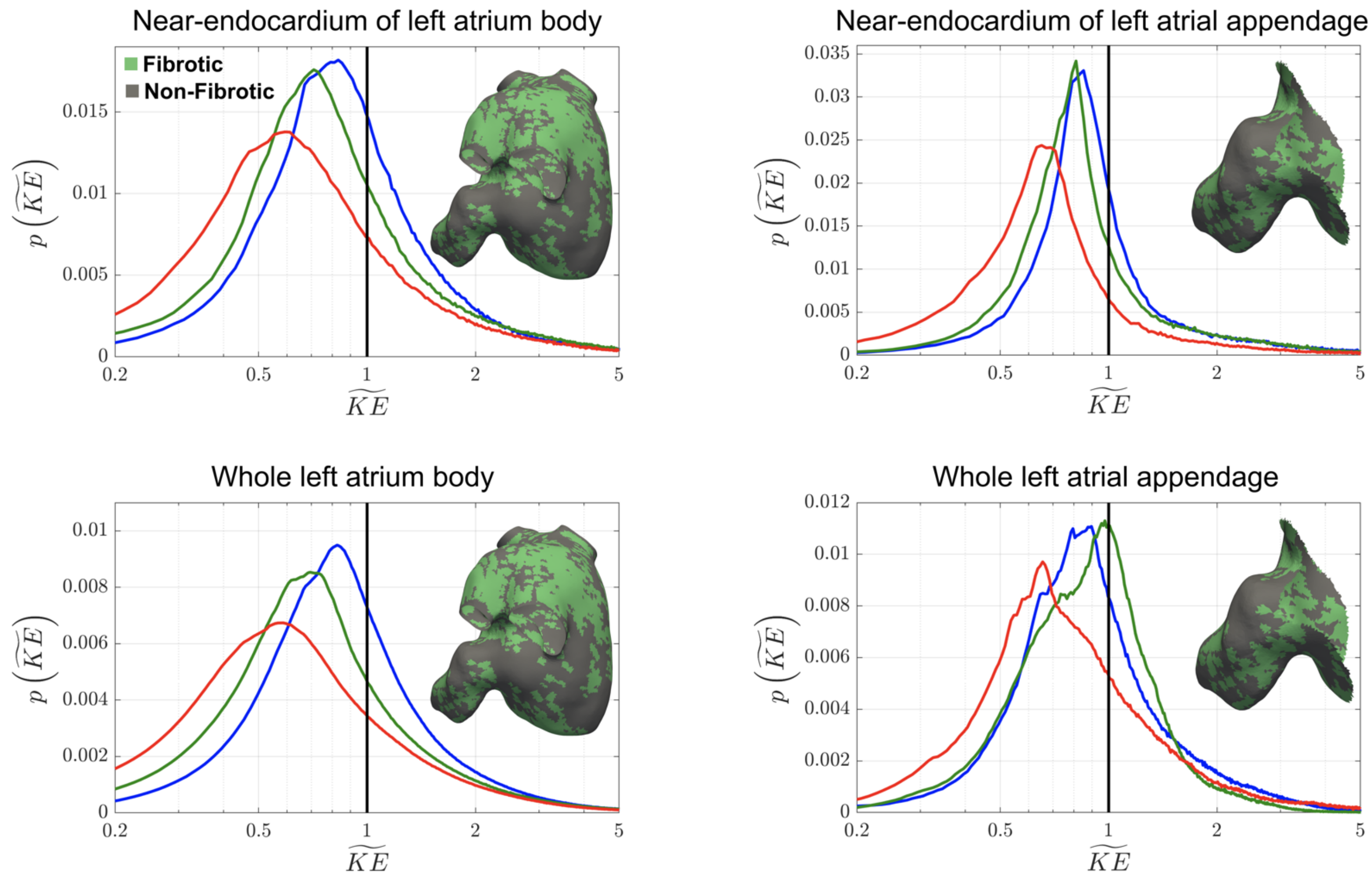


##### *B)* Probability distribution function of normalized kinetic energy ($\widetilde{KE}$) Fib07

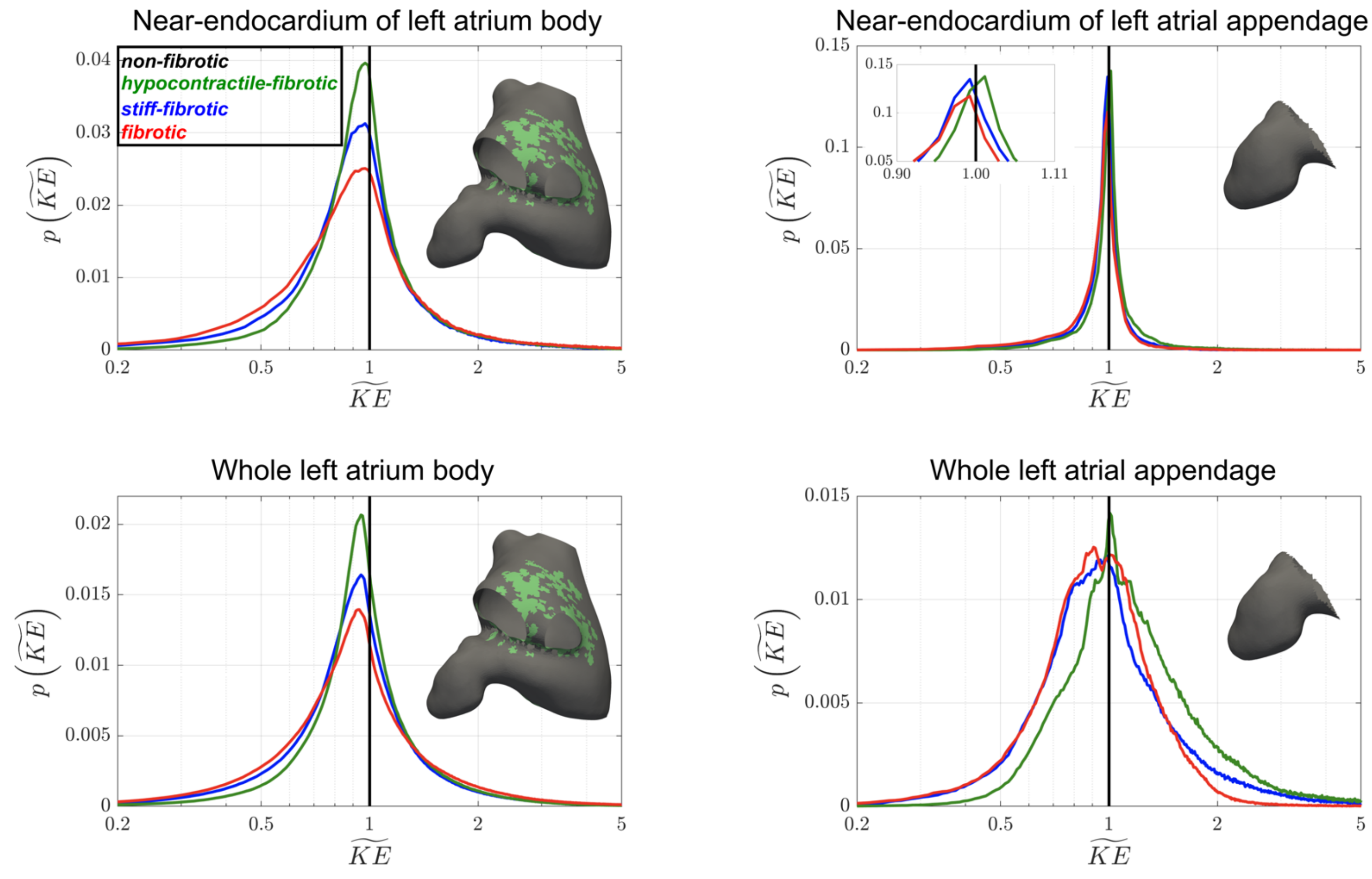

**Figure SI 1. Dissecting the hemodynamic effects of fibrotic tissue stiffening and hypocontractility . A)** Probability distribution function of normalized kinetic energy ($\widetilde{KE}$) in the left atrial body (left column) and in the left atrial appendage (right column) of subject Fib41 over a phase-averaged cycle (contraction and expansion). Near-endocardium results are shown in upper row and whole chamber results in lower row. The black, blue, green, and red lines represent the *non-fibrotic*, *stiff-fibrotic, hypocontractile-fibrotic,* and *fibrotic* mechanical states, respectively. Insets of left atrium (left column) and left atrial appendage (right column) patient-specific models, including fibrotic (green) and non-fibrotic (gray) points, are inserted in the plots to illustrate the fibrotic maps of each subject. **B)** Same as panel **A** for subject Fib07.

### Dissecting the hemodynamic effects of fibrotic tissue stiffening and hypocontractility

##### *A)* Probability distribution function of normalized kinetic energy ($\widetilde{KE}$) Fib47

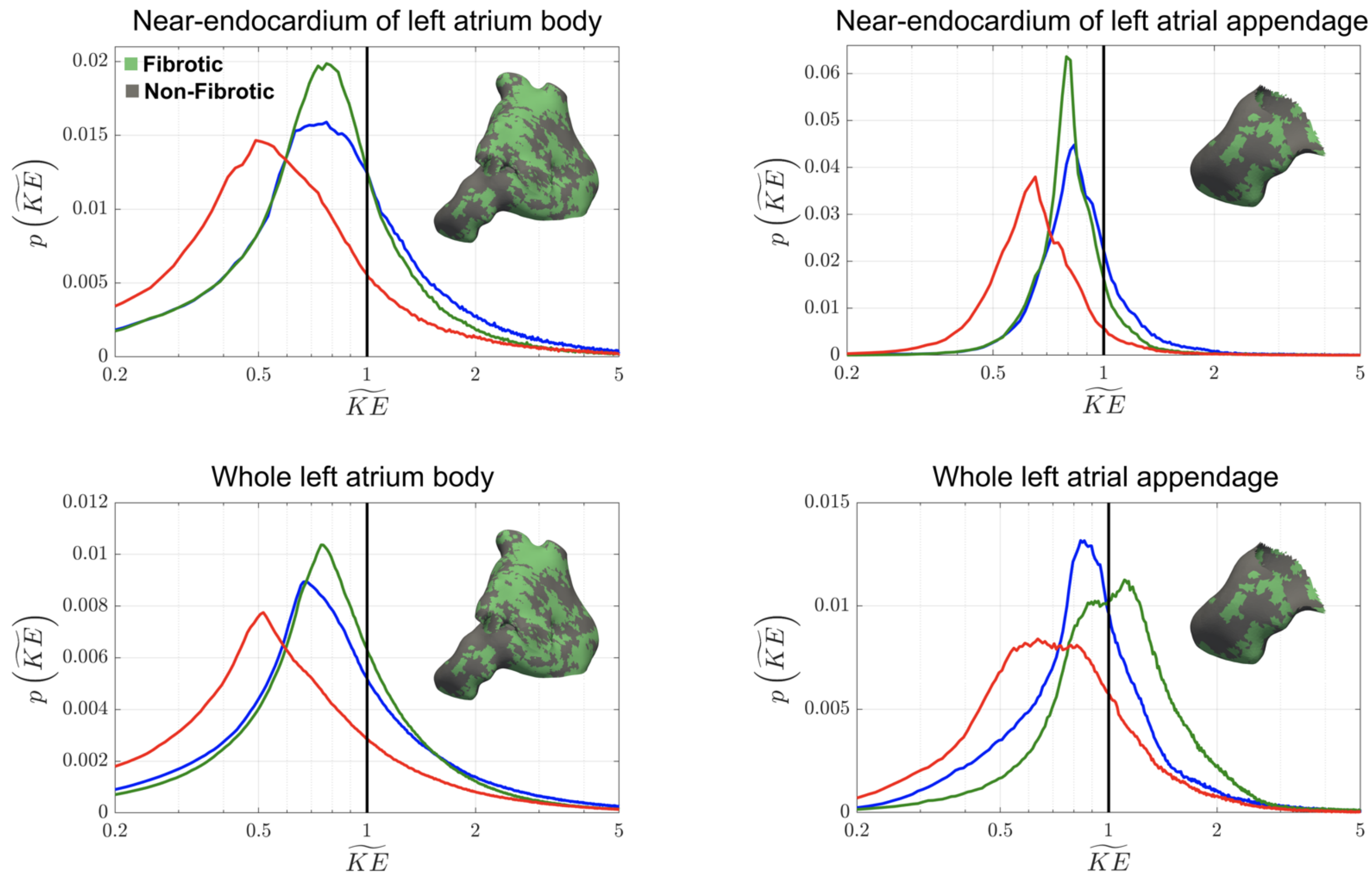


##### *B)* Probability distribution function of normalized kinetic energy ($\widetilde{KE}$) Fib12

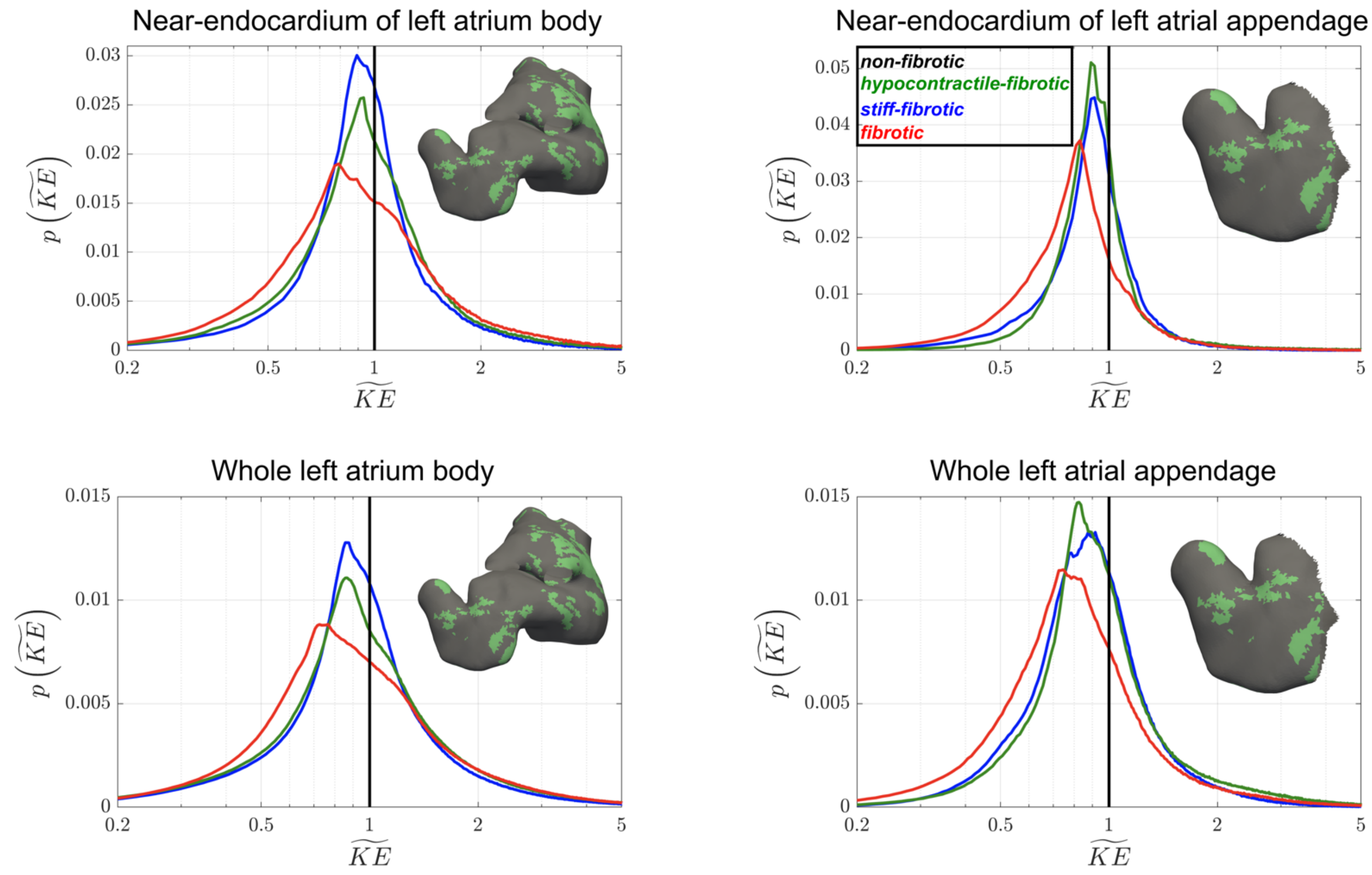

**Figure SI 2. Dissecting the hemodynamic effects of fibrotic tissue stiffening and hypocontractility . A)** Probability distribution function of normalized kinetic energy ($\widetilde{KE}$) in the left atrial body (left column) and in the left atrial appendage (right column) of subject Fib47 over a phase-averaged cycle (contraction and expansion). Near-endocardium results are shown in upper row and whole chamber results in lower row. The black, blue, green, and red lines represent the *non-fibrotic*, *stiff-fibrotic, hypocontractile-fibrotic,* and *fibrotic* mechanical states, respectively. Insets of left atrium (left column) and left atrial appendage (right column) patient-specific models, including fibrotic (green) and non-fibrotic (gray) points, are inserted in the plots to illustrate the fibrotic maps of each subject. **B)** Same as panel **A** for subject Fib12.
